# Filtering cells with high mitochondrial content removes viable metabolically altered malignant cell populations in cancer single-cell studies

**DOI:** 10.1101/2024.10.24.620025

**Authors:** Josephine Yates, Agnieszka Kraft, Valentina Boeva

## Abstract

**Background:** Single-cell transcriptomics has transformed our understanding of cellular diversity in biological systems. However, systematic noise, often introduced by low-quality cells, can obscure biological signals if not properly accounted for. Thus, one of the common quality control steps involves filtering out cells with a high percentage of mitochondrial RNA counts (pctMT), as high pctMT typically indicates cell death. Yet, commonly used filtering thresholds, primarily derived from studies on healthy tissues, may be overly stringent for malignant cells, which often naturally exhibit higher baseline mitochondrial gene expression. We analyzed public single-cell RNA-seq and spatial data to investigate if malignant cells with high pctMT are viable and functionally significant subpopulations.

**Results:** We analyzed nine single-cell RNA-seq datasets from uveal melanoma, breast, lung, kidney, head and neck, prostate, and pancreatic cancers, including 439,507 cells from 151 patients. Malignant cells exhibited significantly higher pctMT than nonmalignant cells without a significant increase in dissociation-induced stress signature scores. Malignant cells with high pctMT showed metabolic dysregulation, including increased xenobiotic metabolism, which is implicated in cancer therapeutic response. Our analysis of pctMT in cancer cell lines uncovered associations with resistance and sensitivity to certain classes of drugs. Additionally, we observed a link between pctMT and malignant cell transcriptional heterogeneity as well as patient clinical features.

**Conclusions:** This study provides a detailed exploration of the functional characteristics of malignant cells with elevated pctMT, challenging current quality control practices in single-cell RNA-seq analyses of tumors. Our findings have the potential to improve data interpretation and refine the biological conclusions of future cancer studies.

## Background

Single-cell transcriptomics studies have led to significant progress in our understanding of tumor biology, paving the way for the development of personalized medicine [1–4]. A crucial early step in processing single-cell RNA-sequencing (scRNA-seq) is implementing rigorous quality control measures to exclude observations that do not represent viable single cells. Following established guidelines [5–8], cells exhibiting a high percentage of mitochondrial RNA content (pctMT) are routinely excluded from the analysis. This practice is based on evidence linking high pctMT to dissociation-induced stress and necrosis [9–11]. However, recent studies have highlighted the limitations of these standard quality control (QC) filters, advocating for novel, data-driven QC metrics [12–15].

Moreover, pctMT has been closely linked to cell-specific metabolic activity, leading to substantial variability across different cell types and often surpassing the thresholds set by traditional filters [8,14,16,17]. For instance, Montserrat-Ayuso and Esteve-Codina [12] argued that conventional mitochondrial filters may inadvertently eliminate healthy cells with high metabolic activity. Additionally, most studies linking pctMT with cell quality have been conducted on healthy rather than diseased tissue, whereas malignant tissues often exhibit higher percentages of mitochondrial counts due to generally elevated mitochondrial DNA (mtDNA) copy number [18] or the activation of the mTOR pathway [19,20]. Hence, using a predefined threshold or median absolute deviations based on the entire cell population to filter out cells with high pctMT in cancer studies might inadvertently remove rare, functionally important subpopulations of malignant cells.

Here, we set out to determine whether subpopulations of malignant cells with high pctMT in cancer indeed correspond to cells suffering from the dissociation-induced stress, empty or broken droplets, or if they represent a viable, functional subpopulation that should be preserved for downstream analysis. By examining publicly available scRNA-seq cancer datasets, we demonstrate that elevated pctMT in malignant cells is largely independent of dissociation-induced stress and that including cells with high pctMT does not significantly compromise dataset quality. We further show that high pctMT malignant cells are metabolically dysregulated and associated with drug response and patient clinical features. Our findings complement current guidelines for processing scRNA-seq datasets and are likely to inform refined quality control strategies in future studies of human cancers.

## Results

### Malignant cells show a significantly higher percentage of mitochondrial RNA than healthy counterparts in samples across cancer types

To determine whether malignant cells exhibit a higher baseline pctMT, we analyzed pctMT levels in both tumor microenvironment (TME) and malignant cells across nine different studies: lung adenocarcinoma (LUAD), small cell lung (SCLC), renal cell (RCC), breast (BRCA), prostate, nasopharyngeal carcinoma (NPC), uveal melanoma, and primary and metastatic pancreatic cancers [4,21–28], spanning the total of 439,507 cells across 151 patients, including 155,573 malignant cells (Fig. 1). We conducted extensive initial quality control (QC) without applying pctMT-based filtering. We evaluated whether this QC approach excluded potential low-quality cells by examining metrics typically associated with cell integrity, as outlined by Ilicic *et al.* [9]. Our analysis confirmed that the cells filtered out by our QC procedure consistently exhibited poor quality metrics despite the QC not explicitly relying on pctMT (Suppl. Fig. S1). Additionally, following recent studies recommending the use of *MALAT1* expression as a QC metric [12,13], we compared the *MALAT1* expression between filtered and retained cells. We found that our filtering process effectively removed cells with high *MALAT1* expression, often associated with nuclear debris, and cells with null *MALAT1* expression, linked with cytosolic debris (Suppl. Fig. S2).

**Figure 1:**
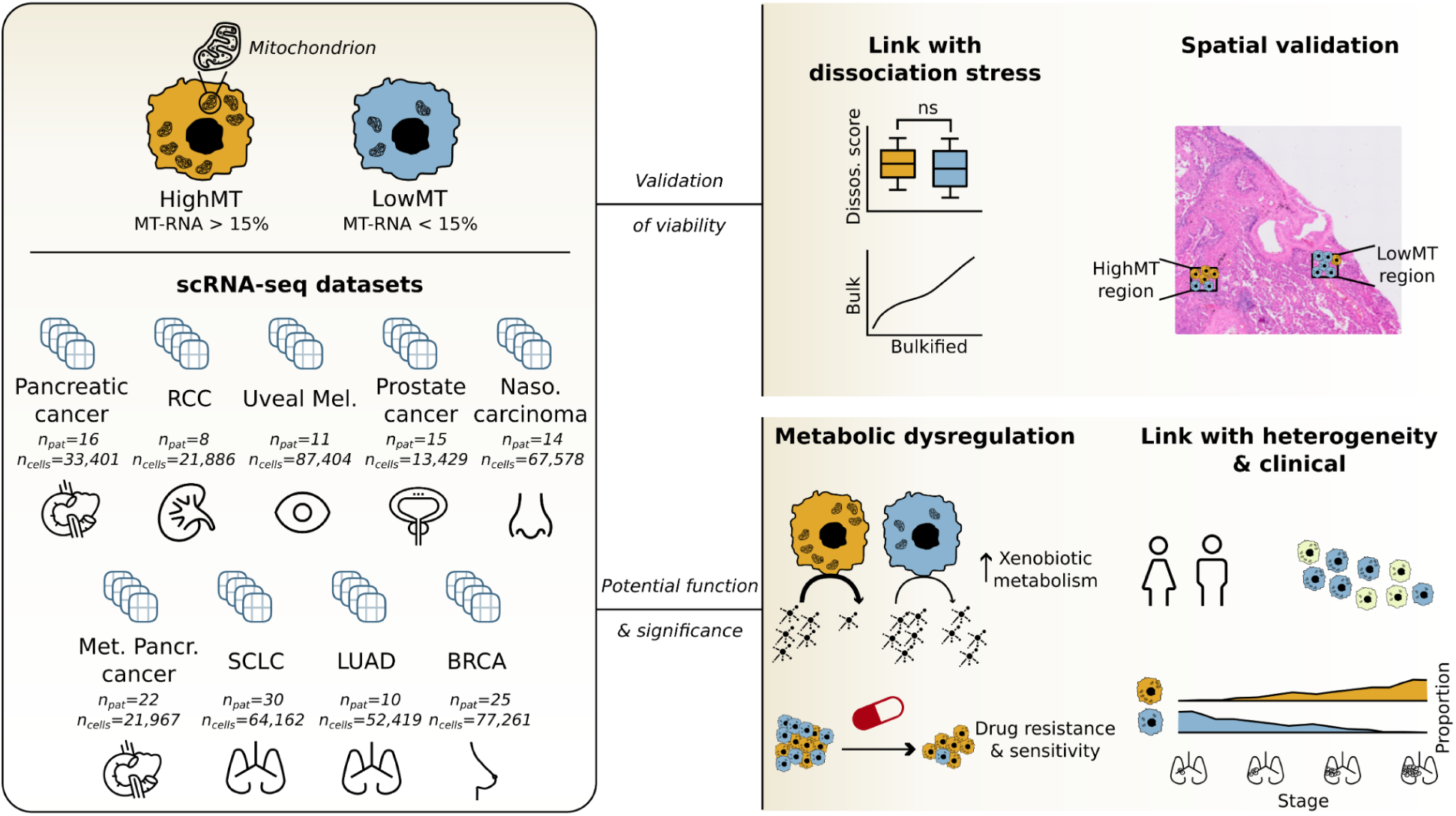
Study Overview. We analyzed nine single-cell cancer datasets [4,21–28] across 151 patients and 439,507 cells from various cancer types, categorizing cells by their percentage of mitochondrial-encoded gene RNA counts (pctMT), with cells above 15% designated as high mitochondrial content cells (HighMT). First, we examined potential links between pctMT and common artifacts, including dissociation-induced stress. We then confirmed regions of high-density malignant HighMT cells in Visium HD slides and explored metabolic dysregulation, notably an increase in xenobiotic metabolism in malignant HighMT cells. We linked cell line pctMT levels to differential drug resistance and sensitivity. Finally, we identified significant associations between pctMT and established cancer cell states, along with key clinical characteristics.

We categorized cells as HighMT or LowMT based on their pctMT values, with those having pctMT above 15% designated as HighMT and those below 15% as LowMT. The value of 15% was chosen as the typical pctMT threshold range used in the non-cancer and cancer studies is 10-20% [24,25,31–35]. We detected significant variability in pctMT distribution between TME and malignant cells across patients, with generally higher median pctMT observed in the malignant cells in both filtered and unfiltered studies (Fig. 2a-b). Overall, 70% of samples (78 out of 111 patients used in this analysis, Methods) had significantly higher pctMT in the malignant compartment (two-sided Mann-Whitney U test p-value < 0.05, Fig. 2a-b). Moreover, across studies of all cancer types, 10% to 50% of tumor samples exhibited a twice higher proportion of HighMT cells in the malignant compartment than in the TME (Methods), indicating a widespread presence of malignant cell subpopulations that would typically be filtered out when the standard 15% cut-off on pctMT is used (Fig. 2a-b). The observed increase in pctMT in carcinomas could be partially explained by the natural variability in pctMT across cell types. Indeed, the basal pctMT of epithelial cells was generally higher than that of other TME components in most cancer types (Suppl. Fig. S3-S11). However, in the majority of cases, the pctMT in the malignant compartment exceeded that of healthy epithelial cells (Suppl. Fig. S3-S11).

**Figure 2:**
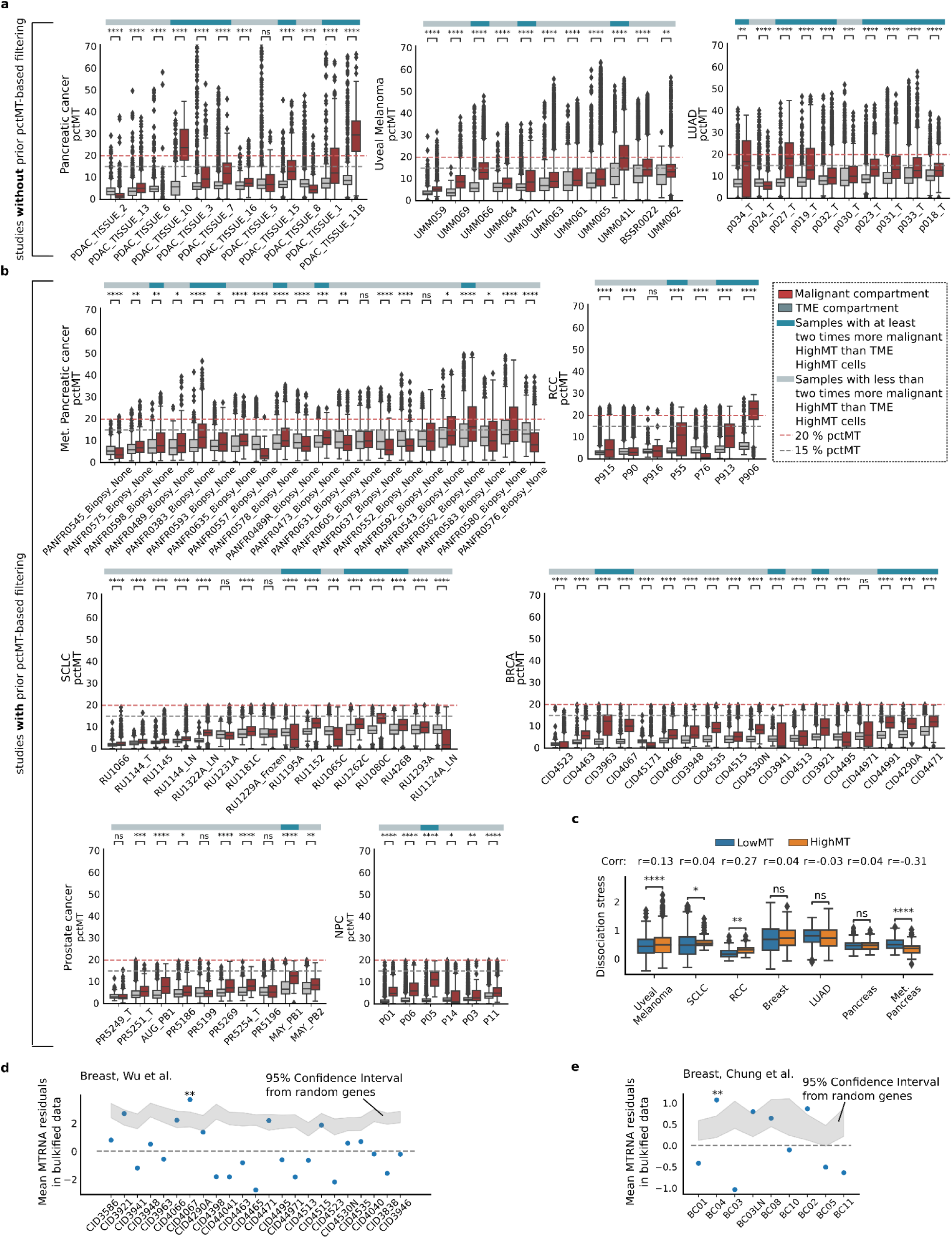
The malignant compartment of multiple cancer types shows subpopulations with high mitochondrial-encoded RNA content. **a-b**, Comparison of mitochondrial RNA percentage (pctMT) between tumor microenvironment (TME) and malignant cells across 111 patients in **a,** unfiltered cohorts and **b,** cohorts with prior pctMT filtering in original studies (Methods) [4,21–26,28]. Patients with too few TME or malignant cells are discarded for this analysis. Patients with more than double the proportion of HighMT malignant cells (pctMT > 15%) compared to TME and with over 15% of HighMT malignant cells are highlighted (blue bar above boxplots). **c,** Distribution of the dissociation-induced stress scores estimated in HighMT and LowMT malignant metacells (pctMT < 15%) across the seven studies selected for the analysis (studies with at least two samples with at least 30% of HighMT malignant metacells). A dissociation stress meta-signature is defined using the common genes in three different dissociation stress signatures [10,11,29]. The point biserial correlation coefficient between the score and HighMT/LowMT status is indicated over the boxplots. **d-e,** Mean of the residuals between the experimental and predicted expression of the 13 MT-encoded protein-coding genes for the paired bulk and single-cell data from the **d,** Wu *et al.* [25] and **e,** Chung *et al.* [30] cohorts. The relationship between bulk and bulkified gene expression is modeled by a polynomial regression. Residuals are computed as the difference between the ground-truth and the predicted bulkified expression. We use an empirical sampling scheme where we compare the mean residuals to that of randomly sampled genes (Methods). The 95% confidence interval of the mean residuals of randomly sampled genes is represented as the shaded gray area, and significance is reported based on Bonferroni-corrected p-values. RCC: renal cell carcinoma; SCLC: small cell lung cancer; NPC: nasopharyngeal carcinoma; LUAD: lung adenocarcinoma; BRCA: breast cancer; Met. Pancr. cancer: metastatic pancreatic cancer; TME: tumor microenvironment. Significance for a-c is computed with a Mann-Whitney U test. ns: 𝑝 > 0. 05; *: 0. 01 < 𝑝 ≤ 0. 05; **: 0. 001 < 𝑝 ≤ 0. 01; ***: 0. 0001 < 𝑝 ≤ 0. 001; ****: 𝑝 ≤ 0. 0001.

### Malignant cells with high mitochondrial content do not strongly express markers of the dissociation-induced stress

We investigated the common hypothesis that the presence of malignant cells with high pctMT in scRNA-seq datasets is due to tissue dissociation protocol inducing cell stress. Utilizing dissociation-induced stress signatures derived from studies by O’Flanagan *et al.*, Machado *et al.*, and van den Brink *et al.* [10,11,29], we constructed a meta score based on genes found across all studies.

To determine whether the HighMT cells in the malignant compartment were associated with dissociation-induced stress without inflating the estimates of statistical significance, we computed metacell expression vectors for each study and excluded studies with insufficient number of HighMT metacells (<30% of all metacells) [36]. In the remaining seven studies, we compared the meta dissociation-induced stress scores between HighMT and LowMT metacells in both healthy and malignant compartments. The results revealed inconsistent patterns: one study indicated lower dissociation-induced stress in malignant HighMT cells, three showed no significant difference, and three showed higher dissociation stress in highMT cells (Fig. 2c). This variability persisted when scoring on a patient-specific basis (Suppl. Fig. S3-S11). Notably, even in the studies where scores of the dissociation-induced stress were higher in the HighMT population of malignant cells, the effect size was small (maximum point biserial coefficient across studies < 0.3), suggesting dissociation-induced stress is unlikely to be the main driver of the HighMT subpopulation in the malignant compartment.

To evaluate whether our QC procedure effectively removed cells stressed by tissue dissociation, or whether adding an additional pctMT filter would further reduce the presence of cells with high stress signature scores, we compared stress signature scores across three groups of malignant cells: cells filtered out by our in-house QC procedure, cells that would be excluded by a pctMT filter, and remaining cells (Suppl. Fig. S12). Our analysis showed no significant increase in dissociation-induced stress scores among QC-passing HighMT cells, suggesting that the pctMT filter does not affect the proportion of cells with high stress signature scores. Therefore, applying a pctMT filter does not further reduce dissociation-related stress in retained cells.

To further demonstrate that dissociation-induced stress does not strongly drive elevated pctMT in the cancer cells passing other QC measures, we compared mitochondrial gene expression between paired bulk and single-cell RNA-seq datasets from two breast cancer studies [25,30]. Data from the bulk RNA-seq protocol, which does not require a tissue dissociation step, served as a control. We modeled the relationship between bulk and “bulkified” single-cell data and calculated the residuals reflecting the excess of gene expression from mitochondria in the scRNA-seq cells passing QC (Methods). In the Wu *et al.* cohort, only one out of 23 patients showed significantly higher residuals for mitochondrial-encoded genes than random nuclear-encoded genes (FDR-corrected p-value < 0.05); in the Chung *et al.* cohort, one out of nine patients showed significantly higher residuals (Fig. 2d-e). These results, consistent across models (Suppl. Fig. S13), indicate that mitochondria-encoded genes are generally similarly expressed in bulk samples and QC-passing single-cell data, reinforcing the notion that HighMT subpopulations of malignant cells do not primarily arise from dissociation-induced stress.

### Spatial transcriptomics reveals subregions of breast and lung tissue with viable malignant cells expressing high levels of mitochondrial-encoded genes

Despite the fact that we observed weak to no association between pctMT and dissociation-induced stress, we wanted to further exclude the hypothesis of the HighMT cells being necrotic. To determine if HighMT cells represent viable, non-necrotic populations, we examined Visium HD spatial transcriptomics data from one breast ductal carcinoma in situ (DCIS) patient (Fig. 3a-e) and one lung adenocarcinoma (LUAD) patient (Fig. 3f-j; Methods).

**Figure 3:**
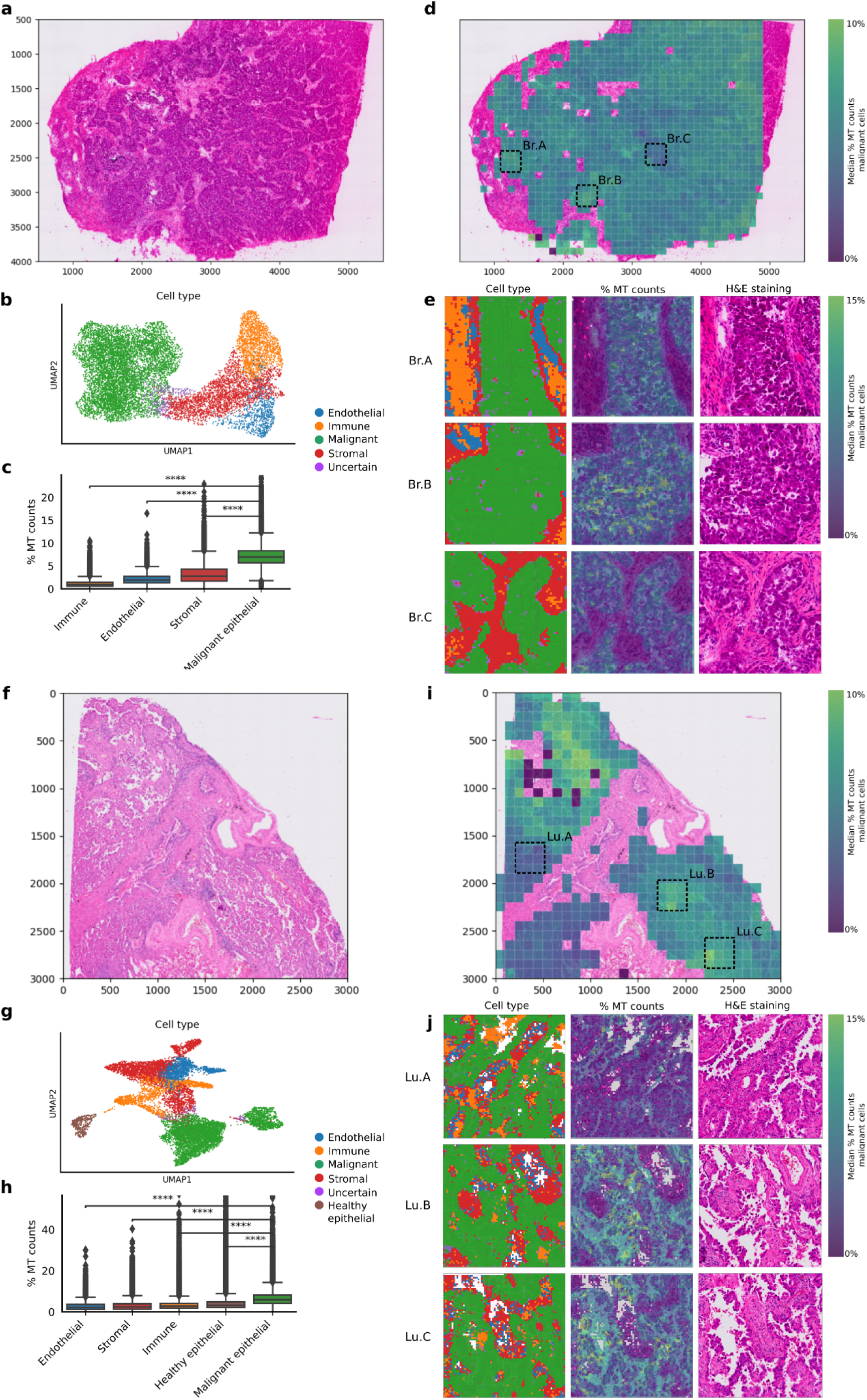
HighMT cells present varied distribution in spatial transcriptomics analyses of lung adenocarcinoma and breast cancer. **a**, H&E staining of breast ductal carcinoma in situ (DCIS) analyzed with Visum HD. **b,** UMAP representation of the “metacells” in the tissue with cell type annotations. We use the Visium 8µm spots (equivalent to single-cell resolution) log1p-normalized gene expression to perform Leiden clustering to define clusters, each aggregated into a “metacell” (Methods). Four primary cell type categories are identified, with copy number variation distinguishing malignant from healthy cells. **c,** Distribution of the pctMT (% MT counts) across cell types in Visium 8µm spots, analyzed by a Mann-Whitney U test (****: *p*<0.0001). **d,** H&E image overlay showing median mitochondrial count percentages of malignant spots in the 100x100px patch. Regions with too few cells are excluded. Breast regions of interest are marked as Br.A, Br.B, and Br.C. **e,** Spot-level cell type annotations, pctMT values, and H&E staining for regions of interest, with cell type annotations derived from metacell data. **f-j,** Same analyses as **a-e** for lung adenocarcinoma (LUAD).

Utilizing transcriptomic data at an 8 µm resolution, equivalent to single-cell resolution as most mammalian cells range from 8-100 µm in size [37], we aggregated spots into metacells and annotated cell types using canonical marker expression and copy number variation analysis in DCIS (Fig. 3b) and LUAD (Fig. 3g). The uncovered copy number variation profiles reflected the published DCIS [38] and LUAD [39] profiles (Suppl. Fig. S14). Consistent with single-cell RNA-seq findings, malignant cells exhibited a significantly higher pctMT than cells in the surrounding tumor microenvironment (TME) in both DCIS and LUAD, with numerous HighMT cells (pctMT > 15%) (Fig. 3c, h).

To assess the spatial distribution of HighMT malignant cells, we calculated the median pctMT across 100x100px patches of malignant spots in both DCIS and LUAD tissues (Fig. 3d, i). This analysis revealed spatial variability, with localized regions showing higher median pctMT among malignant cells. In DCIS, we focused on three regions: Br.A and Br.B (high pctMT) and Br.C (low pctMT) (Fig. 3e). Each region showed consistent malignant cell morphology, but spot-level pctMT varied, with malignant cells displaying significantly elevated pctMT compared to adjacent non-malignant cells. Similarly, in LUAD, regions Lu.B and Lu.C showed markedly higher pctMT than region Lu.A (Fig. 3j).

The findings from spatial transcriptomics confirm that, independent of dissociation stress, malignant cells frequently display elevated pctMT and are variably distributed across tumor regions. This supports the conclusion that viable malignant cells with high pctMT constitute a distinct subpopulation within tumors, observable even without dissociation-induced artifacts.

### Cells with high mitochondrial content have gene signatures associated with mitochondrial transfer and fission

To understand potential mechanisms driving higher pctMT in subpopulations of malignant cells, we explored the link between mitochondrial DNA and RNA content. Previous studies using single-cell and bulk tumor data have shown that transcription of MT-encoded genes positively correlated with the mitochondrial DNA content across healthy and diseased tissues [18,40–43]. Moreover, Kim *et al.* analyzed matched mitochondrial DNA copy number and nuclear DNA data and observed that clones with increased MT-DNA to nuclear DNA ratio (MNR) were associated with higher transcription of mitochondrially encoded oxidative phosphorylation (OXPHOS) genes [44]. To assess whether a similar association is observed between MNR and pctMT in matching clones, we used available data from three ovarian cancer samples and six engineered hTERT cell lines from Kim *et al.* Overall, we observed a positive association between MNR and pctMT (Suppl. Fig. S15).

Higher MT-DNA can result from several mechanisms, including mitochondrial fission [45] or horizontal mitochondrial transfer between TME and malignant cells [46–49]. We assessed the mitochondrial fission activity and the activity of mitochondrial transfer in malignant cells by scoring metacells with the gene ontology (GO) fission signature (GO:0090140) and a recently derived gene signature describing a cancer cell phenotype linked with receiving mitochondria from T-cells [50]. We observed significantly higher scores of one or both signatures in the HighMT malignant cells compared to LowMT ones in five out of seven studies (Fig. 4a-b), with the strongest effect observed in RCC for fission (point biserial correlation coefficient = 0.54, p-value < 0.001) and SCLC for mitochondria transfer (point biserial correlation coefficient = 0.36, p-value < 0.001). These results indicate that higher fission and/or mitochondria transfer from TME might be the driver of higher MT-DNA content and, as such, of higher MT-RNA expression in HighMT cells.

**Figure 4:**
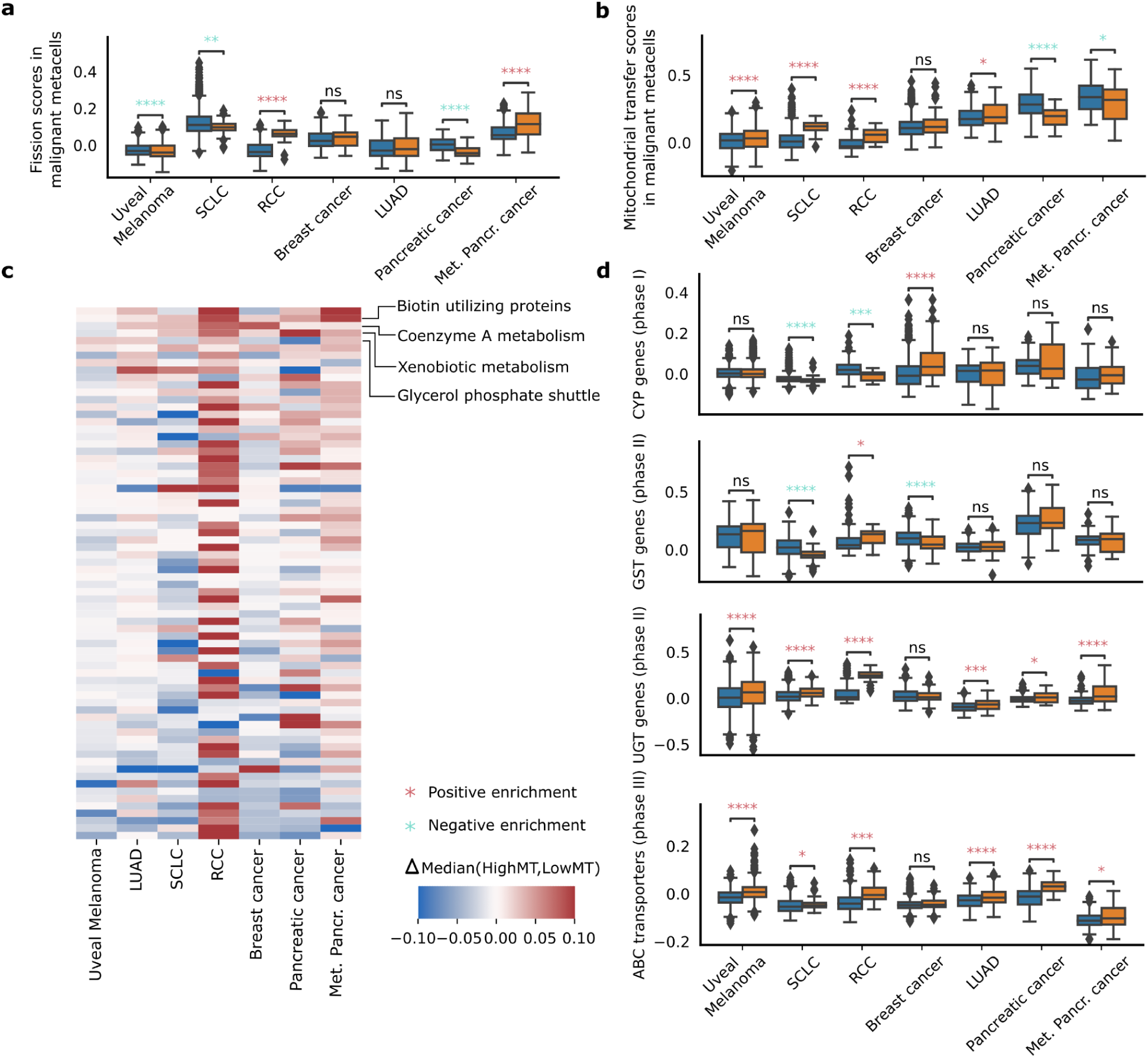
Transcriptomic and metabolic characterization of malignant cells with high mitochondrial content. **a**, Distribution of metacell scores of mitochondrial fission across malignant compartments. **b,** Distribution of metacell scores of mitochondrial transfer across malignant compartments. Significance is computed using a Kruskall-Wallis test. **c,** Heatmap of the dysregulation of the 72 MitoCarta metabolic pathways. The hue represents the difference in median score of the pathway between the HighMT metacells and LowMT metacells. Pathways are ordered according to median difference. **d,** Distribution of signature scores of genes involved in xenobiotic metabolism in the seven studies. The score of CYP genes (phase I), UGT and GST genes (phase II), and ABC transporters (phase III) are compared between HighMT and LowMT metacells for each study. Significance is computed using a Mann-Whitney U test. ns: 𝑝 > 0. 05; *: 0. 01 < 𝑝 ≤ 0. 05; **: 0. 001 < 𝑝 ≤ 0. 01; ***: 0. 0001 < 𝑝 ≤ 0. 001; ****: 𝑝 ≤ 0. 0001.

### Malignant cells with high mitochondrial content present dysregulation of metabolic pathways

Given the essential role of mitochondria in cell metabolism, we hypothesized that malignant cells with high mitochondrial content might exhibit metabolic dysregulation. To investigate this, we examined mitochondrial-related pathways curated by Mitocarta, which includes pathways involving nuclear-encoded proteins or RNAs that translocate to mitochondria [51] (Fig. 4c). We found that the four most consistently upregulated pathways in the studied cancer types were the glycerol phosphate shuttle (7/7), biotin-utilizing proteins (7/7), coenzyme A (CoA) metabolism (6/7), and xenobiotic metabolism (6/7), all with established roles in cancer [52–57]. Notably, oxidative phosphorylation—a core mitochondrial function—was significantly upregulated only in RCC and metastatic pancreatic cancer (Suppl. Table S1), with other cancer types showing no shift or slight downregulation. These results indicate that HighMT cells display notable metabolic dysregulation.

### Cells with high mitochondrial content show increased xenobiotic metabolism through higher drug metabolizing enzymes and ABC transporters

Given our observation of the consistent increase of xenobiotic metabolism gene signature scores in malignant HighMT cells across cancer types, and its implication in cancer therapeutic response [58,59], we further characterized the activity of this pathway by evaluating the expression of genes involved in all three phases of xenobiotic metabolism: phase I cytochrome P450 (CYP) genes, phase II UDP-glycosyltransferase (UGT), and glutathione S-transferase (GST) genes, and phase III ABC transporters (Methods) [60].

We found that HighMT cells showed prominent upregulation of phase II and phase III genes (Fig. 4d). ABC transporters were notably upregulated across all seven studies, reaching statistical significance in six. UGT genes were also significantly elevated in six of the seven datasets. In contrast, phase I genes showed significant upregulation only in the breast cancer dataset. This consistent pattern may reflect the known dependence of ABC transporter-mediated chemoresistance on mitochondrially produced ATP [61].

### Cell lines with higher mitochondrial content show resistance to metabolic drugs and sensitivity to targeting EGFR signaling

Given the high scores of xenobiotic metabolism gene signature in HighMT malignant cells, we further explored the link between the level of expression of mitochondrial RNA and the resistance of cells to commonly used drugs. We analyzed the association between the half-maximal inhibitory concentration (IC50) and mitochondrial content in cell lines from the Cancer Cell Line Encyclopedia (CCLE) [62]. Samples from CCLE showed diverse levels of expression of mitochondrial RNA, ranging from 4% median pctMT in glioblastoma to 14% median pctMT in head and neck squamous cell carcinoma (Suppl. Fig. S16).

We observed a consistent and significant association between elevated pctMT and increased drug resistance, as indicated by higher IC50 values across cell lines with high pctMT (Methods). To confirm the robustness of these associations, we conducted an empirical permutation test, which demonstrated that the observed distribution of correlations significantly diverged from random, particularly in the tails (Suppl. Fig. S16). The top 15 drugs with the highest median resistance across cancer types were significantly enriched in drugs targeting metabolism (Fig. 5a). These included Daporinad, which targets nicotinamide phosphoribosyltransferase (NAMPT) [63], BX-912, which targets PDK1 [64,65], and CAP-232, which targets glycolysis. Many of the other drugs to which cells showed the highest resistance, although not directly associated with metabolism, targeted proteins involved in mitochondrial dynamics. This included MIM1, which targets MCL-1, involved in mitochondrial dynamics [66], MCT4_1422, which targets MCT4, a lactate transporter [67], XMD15-27, which targets CAMK2, linked to mitochondrial-dependent apoptosis [68], and BMS-345541, which targets IKK1, involved in mitochondrial network dynamics [69] (Fig. 5b).

**Figure 5:**
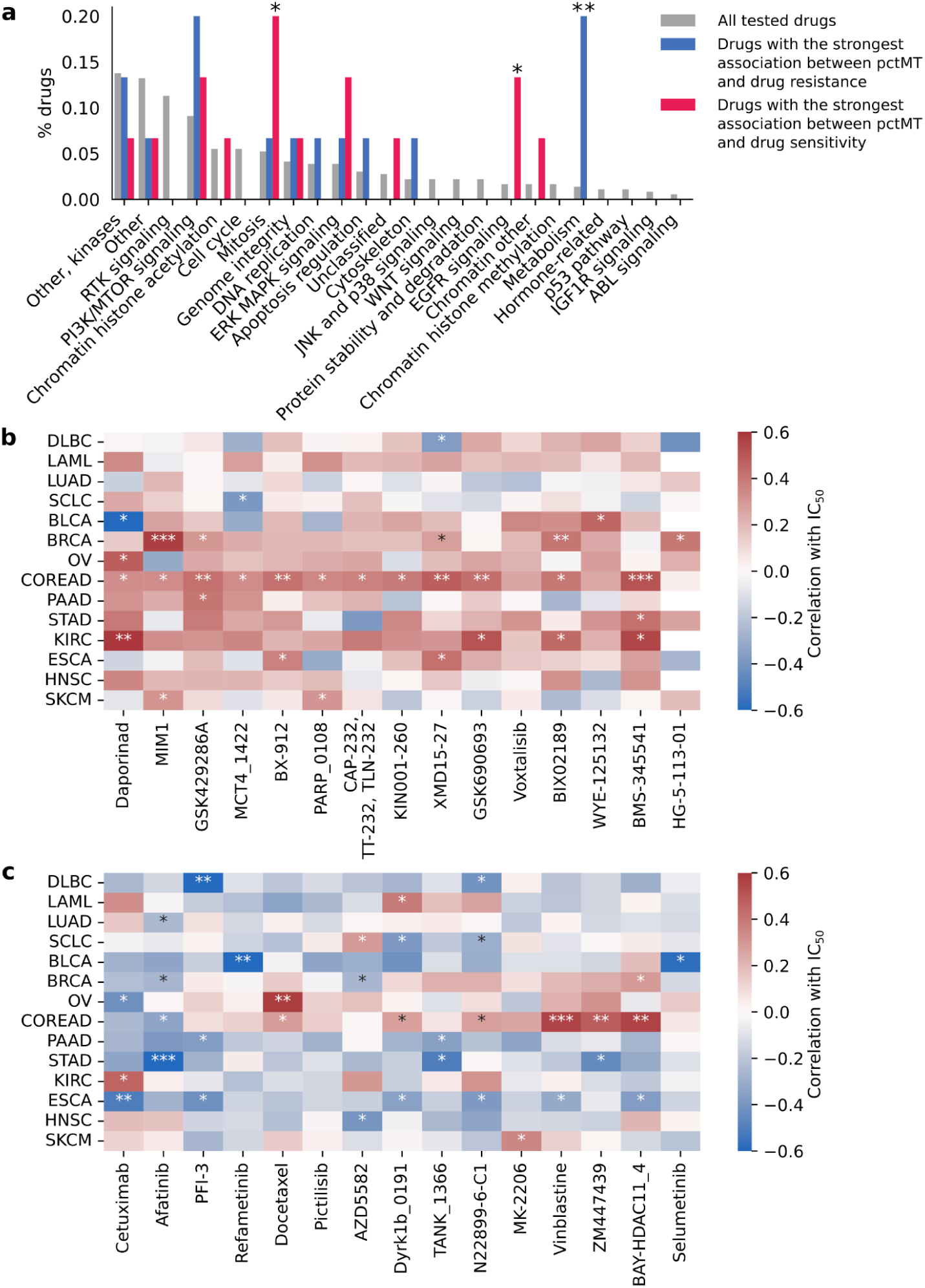
Cell lines with higher mitochondrial content show differential resistance and sensitivities to drugs. **a**, Comparison of the function of the top 15 drugs with the highest association between pctMT and drug resistance (resp. drug sensitivity) and the set of tested drugs. All drugs tested on the CCLE are classified into categories according to their target. The fraction of drugs falling into each category is plotted. Significance is computed using a Fisher exact test. **b-c,** Correlation between the pctMT of cell lines stratified by cancer type and IC50 of specific drugs for the top 15 drugs with the highest median correlation across cancer types (b) and the top 15 drugs with the lowest median correlation across cancer types (c). For each cancer type, Pearson’s correlation between pctMT and IC50 of all cell lines is computed. Significance is computed using Student’s t-test. *: 0. 01 ≤ 𝑝 < 0. 05; **: 0. 001 ≤ 𝑝 < 0. 01; ***: 𝑝 < 0. 001

We also found that higher pctMT in cell lines was consistently linked to higher sensitivity to drugs targeting EGFR signaling or mitosis (Fig. 4a). Specifically, higher pctMT correlated with increased sensitivity to common chemotherapy agents such as Docetaxel and Vinblastine (Fig. 5c). Interestingly, the highMT cells in most cancer types show an increase in expression of EGFR family genes, mostly *ERBB3*, which might partially explain increased sensitivity (Suppl. Fig. S17). Although reports show that EGFR translocates to the mitochondria and is associated with metastasis in lung cancer [70–72], the exact mechanistic link between EGFR, increased mitochondrial content, and its role in carcinogenesis warrants further exploration.

### Subpopulations of malignant cells with higher mitochondrial content are linked to previously reported transcriptional states and patient clinical features

Recent studies across various cancer types revealed the presence of diverse transcriptional profiles of malignant cells within individual tumors and their association with patient treatment outcomes [73–75]. Hence, we investigated whether HighMT cells were enriched in cell populations exhibiting certain previously reported transcriptional programs and states [76,77].

We analyzed gene signature scores characterizing previously reported cancer type-specific transcriptional states in single-cell datasets of SCLC [78], breast [25], uveal melanoma [28], RCC [4], lung adenocarcinoma [79], and pancreatic cancer [80] single-cell studies. Malignant HighMT cells showed significant associations with several reported transcriptional states (Fig. 6a, Suppl. Fig. S18). Specifically, HighMT cells had significantly higher scores for tumor-program 1 (TP1) in RCC, neuroendocrine-like (NE) state in SCLC, mucin-related (TFF1+) and immune-rich (MALAT1+) states in primary and metastatic pancreatic cancer, and TNF-α and hypoxia-related state (GM7) and estrogen response program (GM5) in breast cancer.

**Figure 6:**
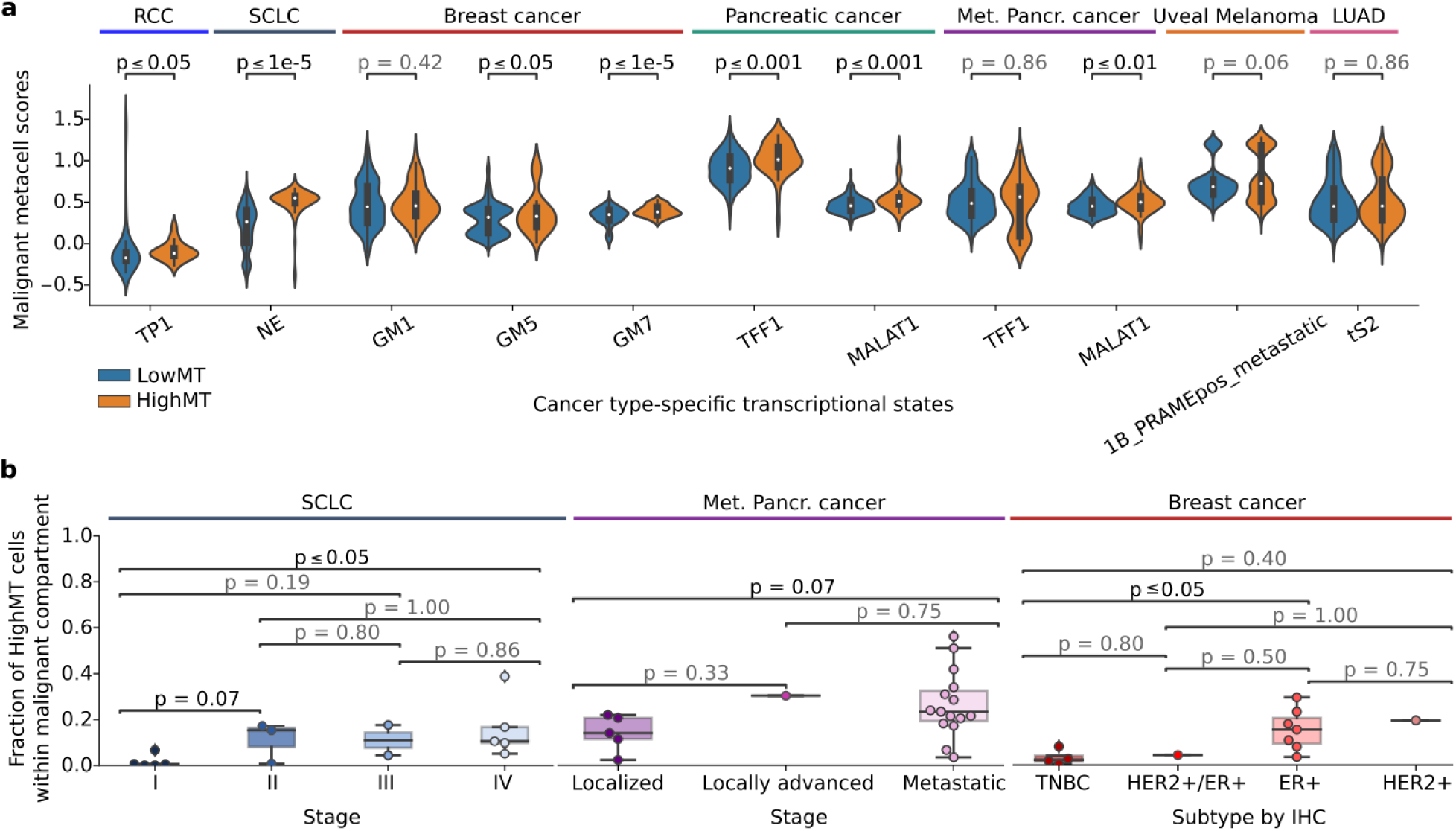
HighMT malignant cells are associated with transcriptional cell states and patient clinical features. **a**, Distribution of scores of previously reported cancer type-specific transcriptional states across HighMT and LowMT cells. The cell states with the median cell-state scores higher in the HighMT than in LowMT cells are shown. Significance is computed using the Mann-Whitney U test on metacells. **b,** Distribution of proportions of HighMT cells within malignant compartment per patient across analyzed datasets and clinical features: stage in SCLC and metastatic pancreatic cancer, and IHC subtype in breast dataset. Significance is computed using the Mann-Whitney U test.

Further, we investigated the link between the proportion of HighMT cells in the malignant compartment and patient clinical features in analyzed single-cell datasets (Fig. 6b). We observed a significant association between the proportion of HighMT malignant cells and stage in SCLC and metastatic pancreatic cancer, with a significantly higher proportion of HighMT malignant cells in more advanced stages (p-value < 0.1). In breast cancer, HighMT malignant cells were significantly enriched in the estrogen receptor-positive (ER+) subtype compared to triple-negative (TNBC) (p-value < 0.05). Taken together, these results show that retaining malignant HighMT cells in scRNA-seq analyses is crucial for accurately capturing tumor heterogeneity and relevant clinical correlations.

## Discussion

Our findings provide evidence that a subpopulation of malignant cells with high mitochondrial content, typically excluded from scRNA-seq analyses, constitutes a metabolically dysregulated and functional subset. High percentages of mitochondrial-encoded gene counts have previously been linked to poor-quality cells, such as damaged droplets or cells affected by dissociation-induced stress [9,10], leading many researchers to filter out cells exceeding a certain pctMT threshold using either static or dynamic criteria. However, recent advancements in data-driven quality control pipelines suggest that setting cell-type specific data-driven QC thresholds can preserve biologically relevant cell populations, such as cardiomyocytes with high pctMT [14]. Our study corroborates these findings by revealing a subpopulation of cancer cells that, when the pctMT filter is relaxed, exhibit dysregulated metabolic functions, notably upregulation of xenobiotic metabolism.

Importantly, our comparison of quality-filtered datasets, with and without pctMT thresholds, shows that including high-quality cells with high pctMT does not affect the overall distribution of dissociation-induced stress scores.

Consistent with previous literature [40], we found that HighMT populations could at least partially be explained by higher MT-DNA content. Higher MT-DNA might be caused by increased mitochondrial fission or horizontal mitochondrial transfer, as previously described [50], and linked to high pctMT in our analysis across several datasets. Hence, the presence of HighMT populations, rather than being caused by poor quality cell capture in the single-cell protocol, can be due to a biologically driven increase in MT-DNA content.

We observed general metabolic dysregulation and upregulated activity of several processes, including xenobiotic metabolism, in HighMT malignant cells. This was mirrored by increased resistance to metabolic drugs in cell lines with high pctMT, suggesting clinical relevance in patient stratification and potential new avenues for combined therapies. These results, together with the association between HighMT cells and previously described transcriptional states, the overrepresentation of HighMT cells in patients of specific molecular subtypes, and the correlation between HighMT subpopulations and tumor stage, suggest the significant role of high pctMT malignant populations in cancer and the importance of including them in analyses.

When further investigating the potential function of malignant HighMT cells, we found that these cells exhibited upregulation of phase II and III genes involved in xenobiotic metabolism, with ABC transporters consistently upregulated across multiple studies. The interdependence between ABC transporter-mediated chemoresistance and mitochondrial ATP production, as highlighted in recent studies [61], may explain this consistent association. Given the limited effectiveness of ABC transporter inhibitors in reversing drug resistance in clinical settings [56], combining these inhibitors with mitochondrial inhibitors could be essential for overcoming resistance.

Several limitations should be acknowledged. First, although we used publicly available data, it was difficult to collect comprehensive datasets with no filters applied on pctMT during data preprocessing. Consequently, some of our datasets do not contain cells with very high pctMT, as these cells were prefiltered. Second, while we utilized previously identified dissociation-induced stress signatures to estimate the stress levels in cells with high pctMT, we lacked definitive ground truths (*e.g*., FACS-sorted stressed cells). Thus, our conclusions regarding the dissociation stress-pctMT relationship require further experimental validation. Third, our spatial analysis of co-existing HighMT and LowMT regions was limited to two samples, restricting the generalizability of our findings. Fourth, the signature we used for horizontal mitochondrial transfer was limited to transfer from T-cells and thus did not take into consideration potential horizontal transfer from other TME compartments. Finally, the link between pctMT and drug resistance and sensitivity was mostly conducted on cell lines, warranting further validation.

## Conclusions

This study is the first to establish that in cancer scRNA-seq datasets, malignant cells with high pctMT, usually filtered out by standard QC procedures, are not solely associated with dissociation-induced stress or poor-quality droplets but represent distinct, functional malignant cell subsets with altered metabolic functions and potentially differential drug responses. The inclusion of HighMT cells in cancer studies is crucial for improving the accuracy of patient stratification and identifying novel therapeutic targets. Moving forward, we recommend adopting more lenient or data-driven pctMT thresholds [14,15] to prevent the loss of valuable biological insights that may contribute to advancements in cancer research and treatment.

## Methods

### scRNA-seq preprocessing

We performed stringent quality control on a patient level across the nine included studies: uveal melanoma [28], small cell lung cancer (SCLC) [24], lung adenocarcinoma (LUAD) [27], renal clear cell cancer (RCC) [4], breast cancer (BRCA) [25], prostate cancer [21], nasopharyngeal carcinoma [26], pancreatic [23] and metastatic pancreatic cancer [22]. We followed the standard processing guidelines described at https://www.sc-best-practices.org/preprocessing_visualization/quality_control.html, excluding steps that involved using the percentage of mitochondrial counts as a quality measure. Notably, some studies had already filtered out cells with less than 20% [21,24–26] or 25% mitochondrial counts [4]. Notably, the lung adenocarcinoma dataset contained cell type annotations only for cells kept in their study but provided raw counts for unfiltered cells; we thus assigned cell types using Leiden overclustering and majority voting of the cell types present in the cluster.

First, we removed all cells per patient that were more than 5 median absolute deviations from the median of either the log1p total number of counts in the cell, log1p genes expressed in the cell, or the percentage of counts falling in the top 50 genes. We also excluded cells with fewer than 1,500 total counts, more than 50,000 total counts, and fewer than 500 genes expressed. Then, we identified and removed putative doublets using Scrublet [81].

Next, using the annotated cell types, we inferred copy number variation (CNV) with inferCNV (https://github.com/icbi-lab/infercnvpy). We clustered the cells in CNV space using the Leiden algorithm, assigning clusters a malignant CNV status if more than half of the cells mapping to the cluster were originally annotated as malignant; otherwise, we assigned a non-malignant CNV status. We removed cells with discordant CNV and transcriptomic identity from downstream analyses. For further analysis, we used the counts per 10k transcripts (CP10K) transformation followed by log(1+x) (log1p) transformation.

### Annotating patients with more than double the proportion of HighMT malignant cells compared to TME HighMT cells

For each study, we compared the distribution of the percentage of transcripts mapping to mitochondrial-encoded gene (pctMT) between cells from the tumor microenvironment (TME) and the malignant cell compartment. We assigned cells to a high mitochondrial content status (HighMT) if they presented >15% pctMT; otherwise, we considered them low mitochondrial content (LowMT). We compared the odds ratio of HighMT cells in the malignant and TME compartments in the rest of the samples using the formula:

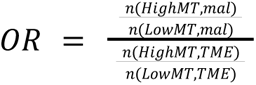

We classified patients as cases if they (*i*) had an OR > 2 and (*ii*) had at least 15% of HighMT cells in the malignant cell compartment; other patients were assigned to controls. For the patient-specific analysis, we removed patients that contained less than 30 malignant or TME cells, and patients that had less than 20 HighMT cells, thus resulting in 111/151 patients. We included only studies comprising more than one case for further analysis.

### Quality metrics and dissociation-induced score computation

The study by Ilicic *et al.* [9] identified seven metrics capable of discriminating between good quality cells and empty/broken cells in a cell-type and technology-agnostic manner, including the Gene Ontology terms *Cytoplasm* (GO:0005737) and *Mitochondrially localized proteins* (GO:0005739), and *mtDNA encoded genes* (equivalent to pctMT) and *Transcriptome variance*. To assess the expression of a gene signature representing GO terms, we applied standard Scanpy scoring [82]. To evaluate transcriptome variance, we calculated the variance per cell using log1p-CP10K-transformed data. We compared these scores between cells filtered out using our quality control (QC) procedure and those retained for downstream analysis.

To construct a dissociation-induced stress score, we aggregated signatures from three external studies. O’Flanagan *et al.* [10] derived a dissociation stress signature from patient-derived breast cancer xenografts, cell lines, and patient cancer cells using 37-degree collagenase dissociation. Machado *et al.* [29] developed a dissociation stress signature based on liver and muscle tissue samples, while Van den Brink *et al.* [11] derived a dissociation stress signature using muscle stem cells. To create a meta-dissociation stress signature, we compiled genes that were consistently found across all three dissociation stress signatures. Cells in our dataset were scored for this meta-dissociation stress signature using standard Scanpy scoring methods.

### Metacell computation

We aggregated single cells of the same type from all 151 patients sequenced through scRNA-seq into metacells to reduce sampling noise and capture underlying transcriptomic distributions, as introduced by Baran et al. [36]. For all remaining seven studies, we implemented metacell aggregation using the Python metacells package (https://github.com/tanaylab/metacells). Metacells were defined as disjoint and homogenous groups of transcriptomic profiles that could potentially arise from the same underlying distribution.

Metacells containing more than 30% of cells with high mitochondrial content were categorized as HighMT metacells, while metacells containing more than 50% malignant cells were classified as malignant. These metacells underwent similar processing as the original scRNA-seq data, including scoring for dissociation stress using the meta-dissociation stress signature applied to log1p-CP10K transformed data. This approach allowed us to analyze and compare transcriptomic profiles at a more aggregated level, focusing on groups that potentially share similar biological characteristics.

### Bulk *versus* bulkified analysis

DNA library preparation for bulk RNA-seq does not include a tissue dissociation step. Therefore, we compared the expression of mitochondrially encoded (MT-encoded) genes between paired bulk RNA-seq and single-cell RNA-seq datasets to assess the potential effects of dissociation-induced stress on MT-encoded gene expression. Specifically, we utilized two datasets with paired single-cell and bulk data: the breast cancer datasets from Wu et al. [25], sequenced using 10X technology, and Chung et al. [30], sequenced using Smart-seq2.

The Wu *et al.* dataset underwent processing using our standard pipeline, while the Chung *et al.* dataset, due to its low cell count per patient, was analyzed collectively rather than on a per-patient basis. We used the Fragments Per Kilobase of transcript per Million mapped reads (FPKM) measure of gene-length corrected gene expression for bulk data from Wu *et al.* dataset; for Chung *et al.* we used the provided Transcript per million (TPM) estimates.

We performed bulkification, *i.e.,* aggregating single-cell measurements into one vector of gene expression per patient to mimic bulk data. Given Smart-seq2 is not naturally gene-length corrected as 10X measurements are, we used the TPM transformation for Smart-seq2 data while we used raw counts for 10X. For Wu et al., we summed raw counts across cells per patient followed by log1p normalization, while for Chung et al., we computed the mean TPM expression across cells per patient.

Due to inherent differences in noise and dropout rates between single-cell and bulk data, direct comparison of bulk and bulkified data is challenging. To model their relationship, we employed polynomial regression, varying degrees from 1 to 6 and evaluating the coefficient of determination (R2) for each. We selected the optimal model complexity based on the elbow of the R2 curve, where further increases in degree yielded minimal R2 improvement.

To assess similarity in MT-encoded gene expression between bulk and bulkified data, we trained a model excluding MT-encoded genes and computed residuals of predicted *vs.* observed bulkified expression for MT-encoded genes. Given their consistent high expression, MT-encoded genes often resulted in higher residuals, potentially affecting model fit. To statistically evaluate these residuals, we performed an empirical test. We randomly sampled genes from the top 500 most expressed genes in each dataset 500 times, trained models on the remaining genes, and computed residuals for these random genes. We calculated one-sided p-values based on how frequently residuals for these random genes exceeded those for MT-encoded genes, setting significance at 0.05.

This methodology allowed us to robustly compare MT-encoded gene expression profiles between bulk and bulkified data, providing insights into potential impacts of dissociation stress on transcriptomic measurements in single-cell RNA-seq studies.

### Spatial transcriptomics Visium HD processing and analysis

For data acquisition, we downloaded two Visium HD samples from the 10X Genomics website: a fresh frozen sample from a patient with breast ductal carcinoma in situ (DCIS) and a Formalin-Fixed Paraffin-Embedded (FFPE) sample from a lung adenocarcinoma (LUAD) patient. To approximate single-cell expression, we utilized the 8µm spot data, as spot size is comparable to the diameter of mammalian cells (8–100 µm). Spots containing fewer than 50 transcripts and genes expressed in fewer than 100 cells were excluded. Counts per 10k (CP10k) normalization was applied, followed by log1p normalization.

Given the sparse and highly correlated nature of Visium HD measurements at the single-spot level, we conducted the analysis in terms of "metacells," or clusters of spatially redundant spots representing aggregated cellular measurements. To construct metacells, we applied Leiden clustering to the 15-nearest neighbor graph, leading to 13,287 and 11,413 metacells in DCIS and LUAD, respectively. These metacells underwent the same CP10k log1p normalization and Leiden clustering as individual spots.

We used canonical marker scoring via Scanpy to assign cell types to each metacell. For LUAD, marker genes were based on major lung compartments from a recent lung cell atlas [31]: epithelial markers (*FXYD3, EPCAM, ELF3*), endothelial (*CLDN5, ECSCR, CLEC14A*), immune (*CD53, PTPRC, CORO1A*), stromal (*COL1A2, DCN, MFAP4*), and neuroendocrine (*CELF3, SLC6A17, CDK5R2*). For DCIS, we used markers from a recent single-cell study [83] identifying epithelial (*EPCAM, KRT7, KRT8*), immune (*CD3D, CD3E, CD79A, CD79B, CD19, MS4A1, CD3G, JCHAIN, MZB1, LYZ, CD68, FCGR3A*), endothelial (*PECAM1, VWF, CLDN5, CDH5, FLT1, RAMP2*), and stromal (*COL1A1, DCN, COL1A2, C1R, ACTA2*) compartments. Cell type assignment within clusters was based on maximum average scoring.

To profile copy number variation (CNV), we applied inferCNV (https://infercnvpy.readthedocs.io/en/latest/index.html), using presumed non-malignant metacells as the reference. Metacells were clustered by CNV profile, and each cluster was categorized as malignant or healthy based on average CNV scores. Final annotations were refined such that healthy CNV epithelial cells were labeled as “healthy” in LUAD and “uncertain” in DCIS, while TME cells with malignant CNV profiles were marked as “uncertain.”

Cell types within each spot were assigned based on the corresponding metacell annotation. We compared pctMT medians between malignant and TME cell types using a Mann-Whitney U test. The spatial distribution of pctMT in malignant cells was assessed by computing median pctMT in 100x100px regions; regions containing fewer than 100 high-quality spots or fewer than 25 malignant high-quality spots were excluded from further analysis.

### Association of pctMT with mitochondrial DNA content

To investigate whether the pctMT was linked to the mitochondrial DNA (mtDNA) content in single-cell data, we used matched single-cell RNA and WES data from Kim et al. [84]. The mtDNA content was evaluated using mtDNA to nuclear DNA ratio (MNR), *i.e*., the number of mtDNA copies per average haploid nuclear genome. Using the clone annotations called by authors, we compared the distribution of pctMT in clones with their distribution of MNR.

### Mitochondrial transfer and fission

We investigated the hypothesis that higher mitochondrial content in cancer cells may be attributed to horizontal mitochondrial transfer from cells within the tumor microenvironment (TME), as suggested by several studies [85–87]. To quantify the extent of mitochondrial transfer, we employed a signature derived from Zhang et al. [88], which characterizes mitochondrial transfer events. Similarly, to evaluate mitochondrial fission, we used the Gene Ontology GO:0090140 gene signature (https://geneontology.org/). Metacells from the analyzed datasets were scored using standard Scanpy scoring based on the above signatures.

### Metabolic dysregulation

To evaluate the extent of metabolic dysregulation in cells, we employed mitochondrial-localized metabolic pathways curated in MitoCarta [51], focusing on genes that reside within mitochondria. We calculated pathway scores for metacells using standard Scanpy scoring using the genes involved in the respective MitoCarta pathways and compared median scores between HighMT and LowMT metacells. Each pathway was characterized by the vector representing the difference between the median scores of HighMT and LowMT metacells. Hierarchical clustering was performed on these pathway vectors across different cancer types using Ward linkage based on Euclidean distances.

Furthermore, to assess the activation of xenobiotic metabolism, we examined genes involved in three phases of this process: phase I enzymes, predominantly cytochrome P450 enzymes involved in oxidation; phase II enzymes, which conjugate phase I metabolites with molecules like glutathione and sulfate to produce hydrophilic compounds; and phase III proteins, primarily ABC transporters facilitating the transport of drugs across cellular membranes [60]. We compared the expression levels of these genes between HighMT and LowMT metacells across all included studies.

### Link between pctMT and drug resistance in cell lines

To investigate the association between higher pctMT and drug resistance or sensitivity, we used paired RNA-seq and drug sensitivity data from the Cancer Cell Line Encyclopedia (CCLE) [62]. First, we extracted raw RNA-seq counts to calculate pctMT for each cell line. Then, we evaluated the correlation between pctMT and the half-maximal inhibitory concentration (IC50) values of all drugs across the dataset for each cancer type. The median correlation across cell lines within each cancer type was computed, identifying the top 15 drugs with the highest and lowest median correlations as the most resistant and most sensitive drugs, respectively.

Drugs were categorized based on their target disruptions; we compared the distribution of these categories between the full set of drugs tested in CCLE and the most resistant or sensitive drugs using the Fisher exact test. This analysis allowed us to evaluate whether specific categories of drug targets were disproportionately represented among the identified resistant or sensitive drugs across cancer types.

### Link between pctMT and previously reported transcriptional cell states

We assess the association between pctMT in malignant cells and expression of cancer type-specific transcriptional states by scoring the expression of respective gene signatures. The signatures were scored using standard Scanpy scoring in metacells and the difference in score distributions between LowMT and HighMT malignant cells was calculated using the Mann-Whitney U test.

### Link between pctMT and clinical information in analyzed single-cell studies

To assess the association between the prevalence of HighMT malignant cells and patient clinical features, we calculated a proportion of HighMT cells within the malignant compartment for each patient and associated it with available clinical features reported in the original studies. The difference between the distributions of the proportion of HighMT cells in each clinical category was evaluated using the Mann-Whitney U test.

## Supporting information

All supplementary materials

## Abbreviations

pctMT: percentage of mitochondrial counts
scRNA-: seq single cell RNA sequencing
TME: tumor microenvironment
MT: mitochondria
HighMT / LowMT: high/low percentage of mitochondrial counts
QC: quality control
H&E: hematoxylin and eosin stains
MNR: mitochondrial DNA to nuclear DNA ratio
CoA: coenzyme A
IC50: half-maximal inhibitory concentration
CNV: copy number variation
CP10K: counts per 10k transcripts
GO: gene ontology
FPKM: fragments per kilobase of transcript per million mapped reads
TPM: transcript per million
NPC: nasopharyngeal carcinoma
LUAD: lung adenocarcinoma
BRCA: breast carcinoma
RCC: renal cell carcinoma
SCLC: small cell lung cancer
CCLE: cancer cell line encyclopedia

## Declarations

### Ethics approval

Not applicable.

### Consent for publication

Not applicable.

### Availability of data and materials

The single-cell studies used in this study can be downloaded from:

- The Gene Expression Omnibus (GEO) website: Breast cancer, Wu et al., at GSE176078; Pancreatic ductal adenocarcinoma, Steele et al., at GSE155698; Prostate cancer, Song et al., at GSE176031; Nasopharyngeal carcinoma, Chen et al., at GSE150430; Breast cancer, Chung et al., at GSE75688
- The Broad single-cell portal: Metastatic Pancreatic cancer, Raghavan et al. at https://singlecell.broadinstitute.org/single_cell/study/SCP1644/microenvironment-drives-cell-state-plasticity-and-drug-response-in-pancreatic-cancer; Renal clear cell cancer, Bi et al. at https://singlecell.broadinstitute.org/single_cell/study/SCP1288/tumor-and-immune-reprogramming-during-immunotherapy-in-advanced-renal-cell-carcinoma#study-summary
- The cancer cell atlas (3CA): Small cell lung cancer, Chan et al., at https://www.weizmann.ac.il/sites/3CA/lung; Lung adenocarcinoma, Bischoff et al., https://www.weizmann.ac.il/sites/3CA/lung; Uveal Melanoma, Durante et al., https://www.weizmann.ac.il/sites/3CA/othermodels
- Zenodo: mtDNA-linked single-cell, Kim *et al.*, https://doi.org/10.5281/zenodo.10498240 The bulk data used in this study can be downloaded from:
- The Gene Expression Omnibus (GEO) website: Breast cancer, Wu et al., at GSE176078; Breast cancer, Chung et al., at GSE75688
- The Cancer Cell Line Encyclopedia (CCLE): for the CCLE RNA-seq and drug sensitivity data https://depmap.org/portal/data_page/?tab=allData

The two samples processed with spatial transcriptomics method Visium HD are freely available on the 10X website: DCIS

https://www.10xgenomics.com/datasets/visium-hd-cytassist-gene-expression-human-breast-can cer-fresh-frozen and LUAD at https://www.10xgenomics.com/datasets/visium-hd-cytassist-gene-expression-human-lung-cancer-post-xenium-expt.

### Code availability

The code used to analyze the data and arrive at the conclusions of the study is available at https://github.com/BoevaLab/MTRNA-sc-cancer.

## Author information

Josephine Yates and Agnieszka Kraft contributed equally as first authors.

### Authors’ contributions

JY, AK and VB designed the study. JY and AK performed computational analyses. JY and AK prepared the manuscript. VB, JY, and AK revised the manuscript. VB supervised the study. All authors read and approved the final manuscript.

## Acknowledgments

We thank Federica Sella, Andréanne Gagné, and Mitchell Levesque for their critical feedback on the work. We would like to thank Dr. Kim Minsoo for his help in sharing data from his recent paper for the pctMT to mtDNA analysis.

## Supplementary Information

### Supplementary Tables

**Supplementary Table 1: Expression of oxidative phosphorylation (OXPHOS) in the HighMT versus LowMT population.** OXPHOS is scored in metacells and the median for the HighMT and LowMT population, the difference in medians, and the p-value and FDR-corrected q-value for the difference computed with a Mann-Whitney U test are indicated for each cancer type.

### Supplementary Figures

**Supplementary Figure S1: Distribution of quality metrics of filtered and kept cells using our in-house procedure**. **a-h,** For each study included in the analysis [1–9], we run the QC procedure described in Methods, that uses exhaustive QC that does not include using the percentage of mitochondrial counts (% MT counts). We visualize quality metrics for the cells removed with this procedure (Filtered) and those used for the rest of the analysis (Kept). Metrics include standard QC metrics (%MT counts, log1p(total counts)), dissociation stress (computed with the meta-signature devised using the three dissociation stress signatures [10–12]), and three metrics described in Ilicic et al. as cell-type agnostic measures to remove broken and empty droplets (transcriptome variance, mitochondria-located proteins Gene Ontology GO:0005739 and cytoplasm-located proteins GO:0005737). Significance is tested using a Mann-Whitney U test. *: 0. 01 ≤ 𝑝 < 0. 05; : 0. 0001 ≤ 𝑝 < 0. 001***; \****\*\*\*: 𝑝 < 0. 0001.

**Supplementary Figure S2: MALAT1 expression distribution in filtered and kept cells.** For each study included in the analysis, we compare the distribution of MALAT1 expression in Filtered and Kept cells.

**Supplementary Figure S3: Distribution per patient of the uveal melanoma study** [8]**, with (a) percentage of mitochondrial counts among malignant and TME cells and (b) dissociation stress among malignant and TME cells**. Dissociation stress is measured by the score of the meta-dissociation stress signature devised as the genes commonly found in all signatures of dissociation (Methods).

**Supplementary Figure S4: Distribution per patient of the small cell lung cancer Chan *et al.* study** [1]**, with (a) percentage of mitochondrial counts among malignant and TME cells,(b) percentage of mitochondrial counts with the separation of TME and normal epithelial cells, and (c) dissociation stress among malignant and TME cells**. Dissociation stress is measured by the score of the meta-dissociation stress signature devised as the genes commonly found in all signatures of dissociation (Methods).

**Supplementary Figure S5: Distribution per patient of the pancreas ductal adenocarcinoma Steele *et al.* study** [4]**, with (a) percentage of mitochondrial counts among malignant and TME cells, (b) percentage of mitochondrial counts with the separation of TME and normal epithelial cells, and (c) dissociation stress among malignant and TME cells**. Dissociation stress is measured by the score of the meta-dissociation stress signature devised as the genes commonly found in all signatures of dissociation (Methods).

**Supplementary Figure S6: Distribution per patient of the metastatic pancreas ductal adenocarcinoma Raghavan et al. study** [5], **with (a) percentage of mitochondrial counts among malignant and TME cells, (b) percentage of mitochondrial counts with the separation of TME and normal epithelial cells, and (c) dissociation stress among malignant and TME cells.** Dissociation stress is measured by the score of the meta-dissociation stress signature devised as the genes commonly found in all signatures of dissociation (Methods).

**Supplementary Figure S7: Distribution per patient of the prostate cancer Song *et al*. study** [2]**, with (a) percentage of mitochondrial counts among malignant and TME cells, (b) percentage of mitochondrial counts with the separation of TME and normal epithelial cells, and (c) dissociation stress among malignant and TME cells**. Dissociation stress is measured by the score of the meta-dissociation stress signature devised as the genes commonly found in all signatures of dissociation (Methods).

**Supplementary Figure S8: Distribution per patient of the renal clear cell cancer Bi *et al.* study** [6]**, with (a) percentage of mitochondrial counts among malignant and TME cells, and (b) dissociation stress among malignant and TME cells.** Dissociation stress is measured by the score of the meta-dissociation stress signature devised as the genes commonly found in all signatures of dissociation (Methods).

**Supplementary Figure S9: Distribution per patient of the nasopharyngeal carcinoma Chen *et al.* study** [3]**, with (a) percentage of mitochondrial counts among malignant and TME cells, (b) percentage of mitochondrial counts with the separation of TME and normal epithelial cells, and (c) dissociation stress among malignant and TME cells**. Dissociation stress is measured by the score of the meta-dissociation stress signature devised as the genes commonly found in all signatures of dissociation (Methods).

**Supplementary Figure S10: Distribution per patient of the breast cancer Wu *et al.* study** [7]**, with (a) percentage of mitochondrial counts among malignant and TME cells, (b) percentage of mitochondrial counts with the separation of TME and normal epithelial cells, and (c) dissociation stress among malignant and TME cells**. Dissociation stress is measured by the score of the meta-dissociation stress signature devised as the genes commonly found in all signatures of dissociation (Methods).

**Supplementary Figure S11: Distribution per patient of the lung adenocarcinoma Bischoff *et al.* study** [9]**, with (a) percentage of mitochondrial counts among malignant and TME cells, (b) percentage of mitochondrial counts with the separation of TME and normal epithelial cells, and (c) dissociation stress among malignant and TME cells**. Dissociation stress is measured by the score of the meta-dissociation stress signature devised as the genes commonly found in all signatures of dissociation (Methods).

**Supplementary Figure S12: Dissociation-induced stress signature expression distribution in filtered and kept malignant cells.** For each study included in the analysis, we compute the dissociation stress for cells filtered using our QC procedure, cells kept using our procedure that present >15% MT counts, and all other remaining cells.

**Supplementary Figure S13: Residuals of MT-encoded genes for two breast cancer cohorts across different modelizations for a, the Wu et al.** [7] **and b, the Chung et al.** [13] **cohorts.** For each patient, paired bulk and single-cell measurements are available. We model the bulkified vs bulk relationship using a polynomial regression; we represent results for polynomial regression of degrees 1 through 6. The residuals represent the difference for each MT-encoded gene between the true and predicted bulkified values (Methods). To estimate whether a patient displays significantly differing residuals, we use an empirical one-sided test that tests the amount of times the mean residuals of the MT-encoded genes is higher than that of the randomly sampled genes with a similar level of expression. The 95% confidence interval of the mean residuals of randomly sampled genes is represented as a shaded gray area. **d-e,** Coefficient of determination (R2) for the different degrees for the **d,** Wu et *al.* and **e,** Chung et *al.* cohorts. The selected model is the one at the “elbow”, i.e., when the increased complexity comes at the cost of minimal improvement of R2.

**Supplementary Figure S14**: **Copy number variation (CNV) analysis of metacells in breast ductal carcinoma in situ (DCIS) and lung adenocarcinoma (LUAD). a-b,** CNV profiles of metacells in **a,** DCIS and **b,** LUAD, with cell types assigned based on maximum canonical marker scores and inferCNV analysis using putative healthy cells as a reference. **c-d,** UMAP visualization of metacells with CNV data in **c,** DCIS and **d,** LUAD, colored by Leiden clustering, average CNV score, and putative cell type annotations. CNV clusters are categorized as malignant or healthy based on CNV scores, and cell type annotations are then adjusted accordingly.

**Supplementary Figure S15:** Comparison of distribution between mitochondrial-to-nuclear gene ratio (MNR) and the percentage of mitochondrial counts (% MT counts) in (a) ovarian cancer patient sample and (b) engineered 184-hTERT cell lines from Kim et al. [14] Cells in scRNA-seq are computationally assigned to clones inferred in the DLP+-sequenced population using the TreeAlign algorithm [15]. The MNR is inferred using the genomic data while the % MT counts is computed using the scRNA-seq data. Clones with higher MNR often exhibit higher % MT counts.

**Supplementary Figure S16: Drug resistance in cell lines. a,** Distribution of the % of mitochondrial counts in cell lines in the CCLE. **b,** Distribution of the median correlation across cell lines of specific cancer types between the pctMT and the IC50 of drugs. Significance of the difference between distributions is computed using a Kolgomorov-Smirnov test.

**Supplementary Figure S17: EGFR family gene expression across cancer types.** The log1p(CP10k) expression distribution is compared across HighMT and LowMT groups using a Mann-Whitney U test.

**Supplementary Figure S18: Association between pctMT and malignant cell states. a,** Distribution of scores of previously reported cancer-specific transcriptional states in LowMT and HighMT malignant cells. Significance is computed using Mann-Whitney U test on metacells. *: 0. 01 ≤ 𝑝 < 0. 05; **: 0. 001 ≤ 𝑝 < 0. 01; ***: 𝑝 < 0. 001.

