## Supplementary material for "Filtering cells with high mitochondrial content removes viable metabolically altered malignant cell populations in cancer single-cell studies": All supplementary materials

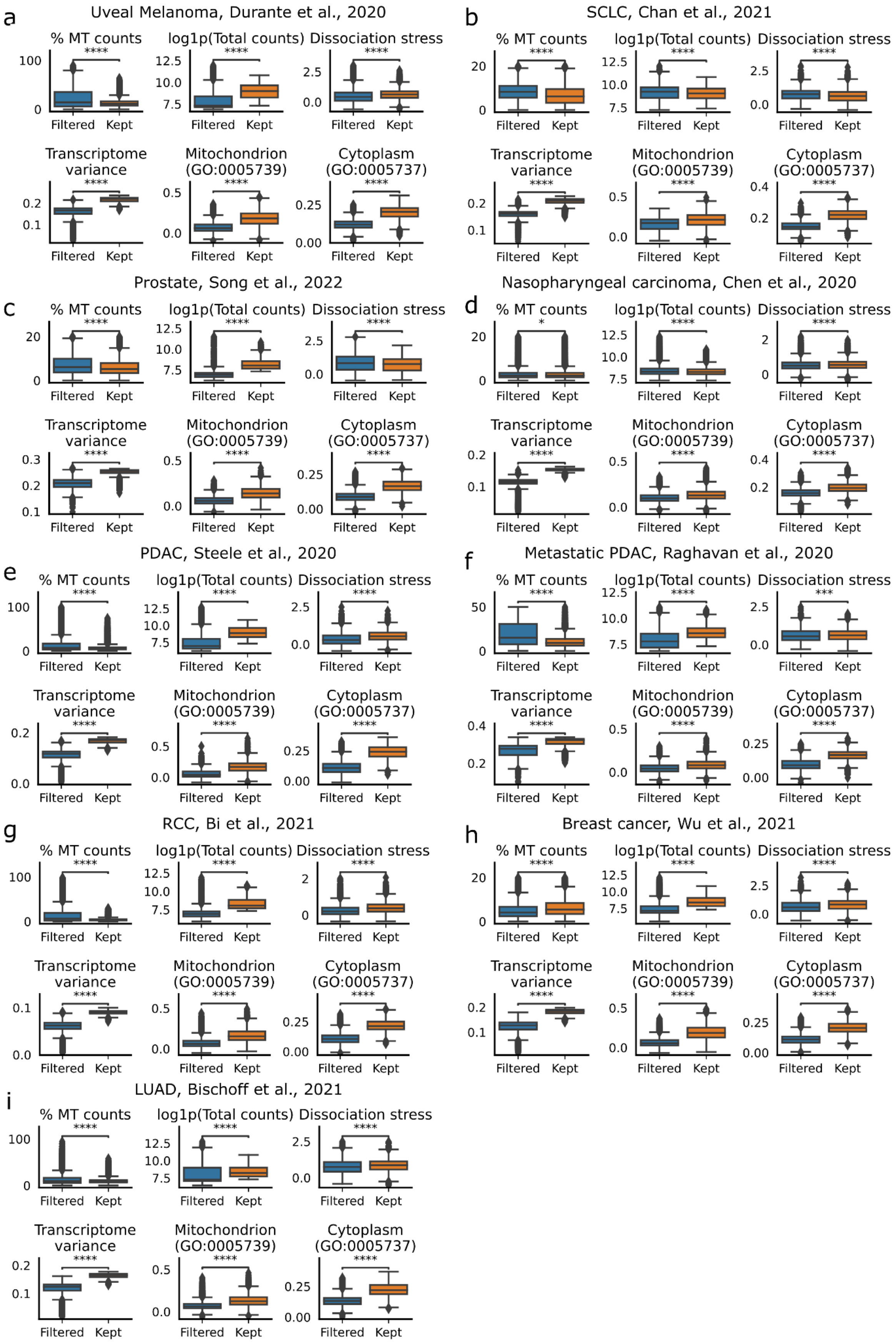
a
Uveal Melanoma, Durante et al., 2020
b
SCLC, Chan et al., 2021
c
Prostate, Song et al., 2022
d
Nasopharyngeal carcinoma, Chen et al., 2020
e
PDAC, Steele et al., 2020
f
Metastatic PDAC, Raghavan et al., 2020
g
RCC, Bi et al., 2021
h
Breast cancer, Wu et al., 2021
i
LUAD, Bischoff et al., 2021
% MT counts
log1p(Total counts)
Dissociation stress
Transcriptome variance
Mitochondrion (GO:0005739)
Cytoplasm (GO:0005737)
Filtered
Kept
****
***
*

**Supplementary Figure S1: Distribution of quality metrics of filtered and kept cells using our in-house procedure**. **a-h,** For each study included in the analysis [1–9], we run the QC procedure described in Methods, that uses exhaustive QC that does not include using the percentage of mitochondrial counts (% MT counts). We visualize quality metrics for the cells removed with this procedure (Filtered) and those used for the rest of the analysis (Kept). Metrics include standard QC metrics (%MT counts, log1p(total counts)), dissociation stress (computed with the meta-signature devised using the three dissociation stress signatures [10–12]), and three metrics described in Ilicic et al. as cell-type agnostic measures to remove broken and empty droplets (transcriptome variance, mitochondria-located proteins Gene Ontology GO:0005739 and cytoplasm-located proteins GO:0005737). Significance is tested using a Mann-Whitney U test. *: $0.01 \leq p < 0.05$; : $0.0001 \leq p < 0.001$; ****: $p < 0.0001$.

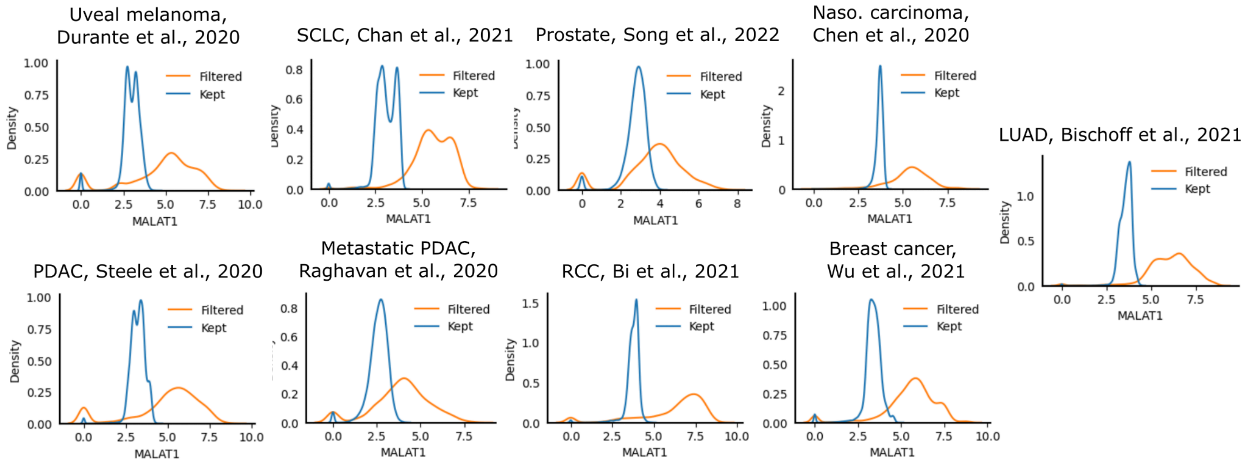


**Supplementary Figure S2: MALAT1 expression distribution in filtered and kept cells.** For each study included in the analysis, we compare the distribution of MALAT1 expression in Filtered and Kept cells.

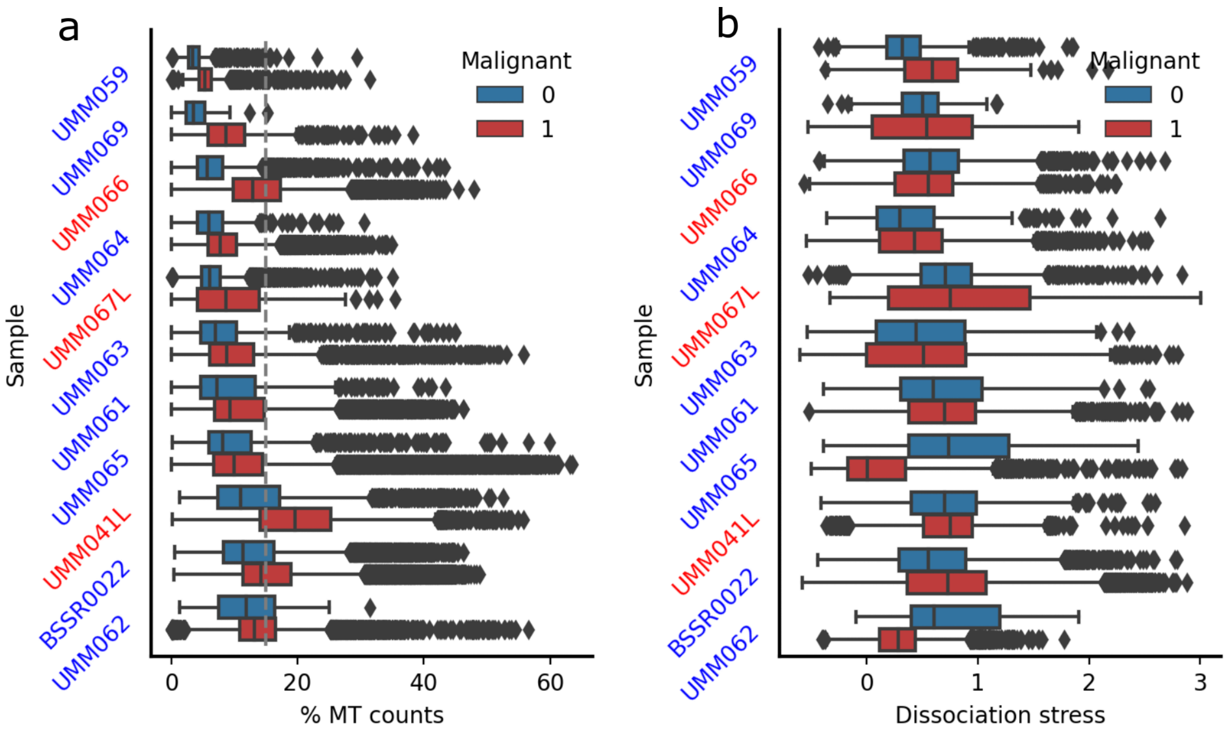


**Supplementary Figure S3: Distribution per patient of the uveal melanoma study** [8]**, with a, percentage of mitochondrial counts among malignant and TME cells and b, dissociation stress among malignant and TME cells**. Dissociation stress is measured by the score of the meta-dissociation stress signature devised as the genes commonly found in all signatures of dissociation (Methods).

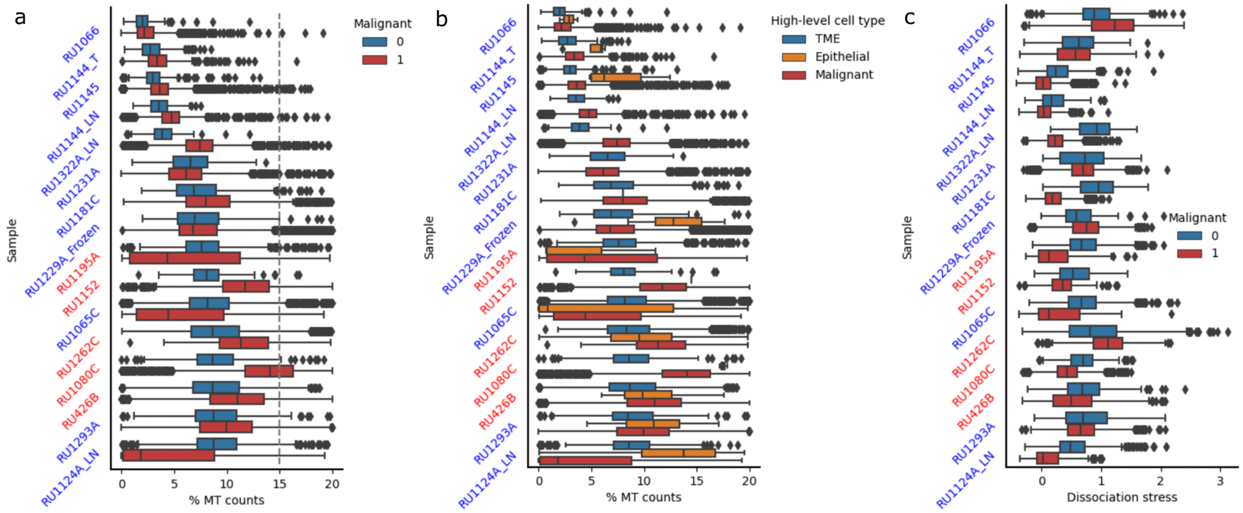


**Supplementary Figure S4: Distribution per patient of the small cell lung cancer Chan *et al.* study** [1]**, with a, percentage of mitochondrial counts among malignant and TME cells, b, percentage of mitochondrial counts with the separation of TME and normal epithelial cells, and c, dissociation stress among malignant and TME cells**. Dissociation stress is measured by the score of the meta-dissociation stress signature devised as the genes commonly found in all signatures of dissociation (Methods).

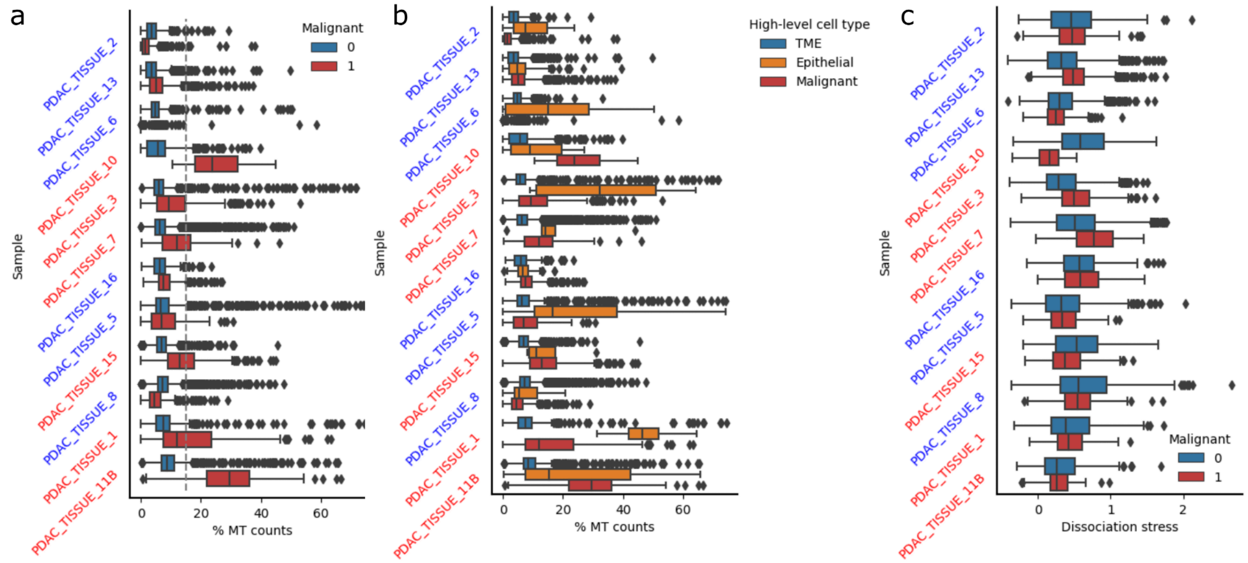


**Supplementary Figure S5: Distribution per patient of the pancreas ductal adenocarcinoma Steele *et al.* study** [4], **with a, percentage of mitochondrial counts among malignant and TME cells, b, percentage of mitochondrial counts with the separation of TME and normal epithelial cells, and c, dissociation stress among malignant and TME cells**. Dissociation stress is measured by the score of the meta-dissociation stress signature devised as the genes commonly found in all signatures of dissociation (Methods).

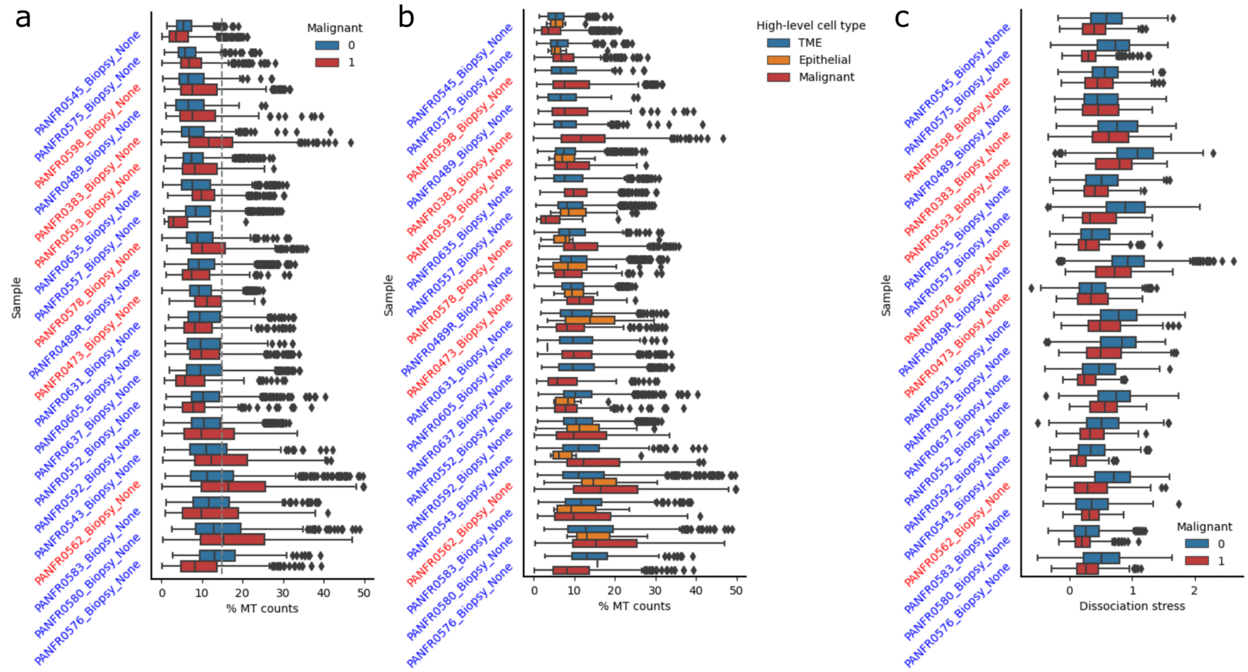

**Supplementary Figure S6: Distribution per patient of the metastatic pancreas ductal adenocarcinoma Raghavan *et al.* study** [5]**, with a, percentage of mitochondrial counts among malignant and TME cells, b, percentage of mitochondrial counts with the separation of TME and normal epithelial cells, and c, dissociation stress among malignant and TME cells**. Dissociation stress is measured by the score of the meta-dissociation stress signature devised as the genes commonly found in all signatures of dissociation (Methods).

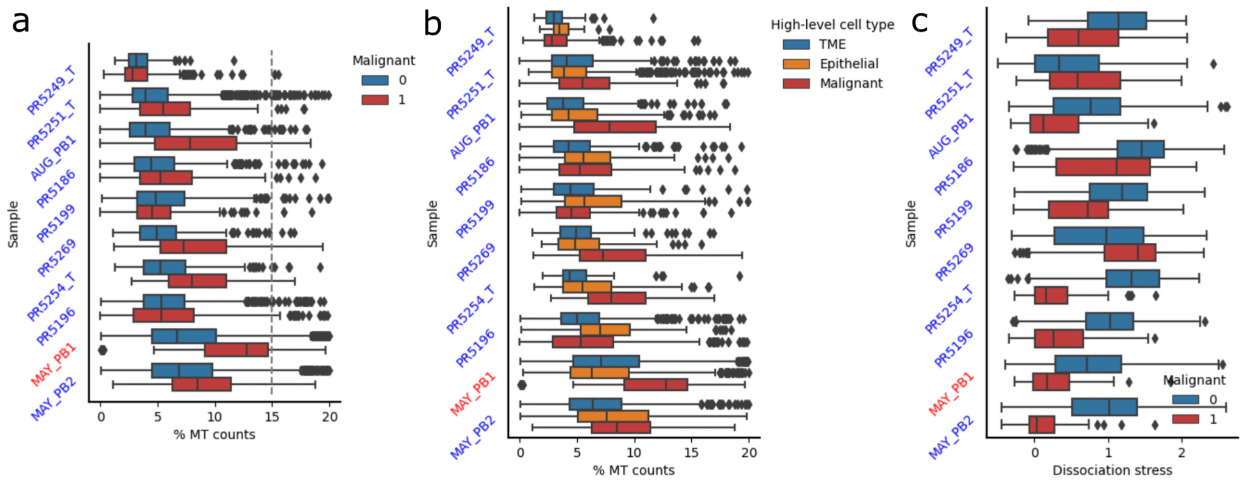


**Supplementary Figure S7: Distribution per patient of the prostate cancer Song *et al.* study** [2]**, with a, percentage of mitochondrial counts among malignant and TME cells, b, percentage of mitochondrial counts with the separation of TME and normal epithelial cells, and c, dissociation stress among malignant and TME cells**. Dissociation stress is measured by the score of the meta-dissociation stress signature devised as the genes commonly found in all signatures of dissociation (Methods).

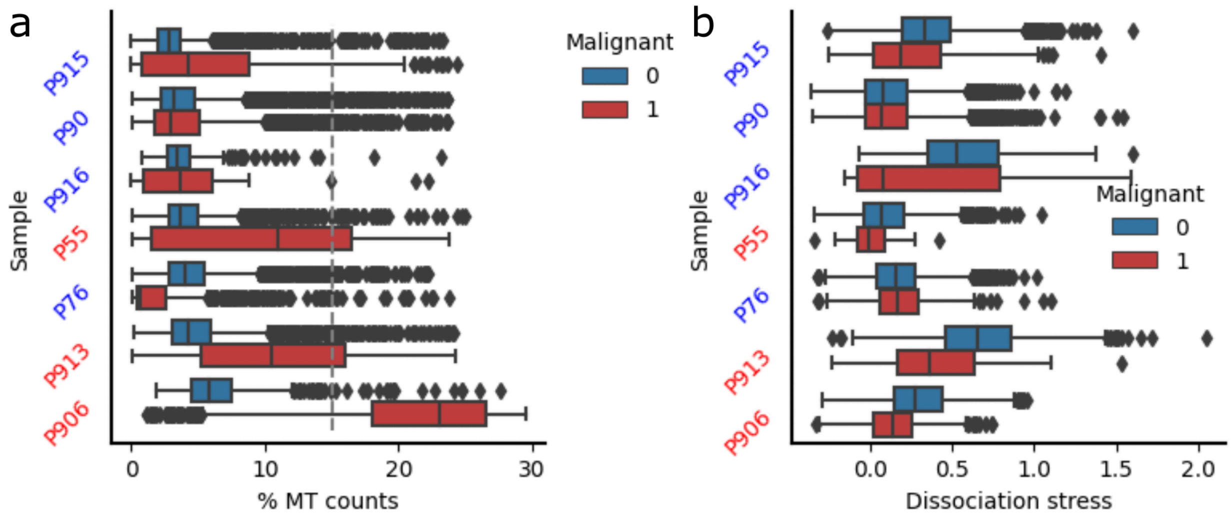


**Supplementary Figure S8: Distribution per patient of the renal clear cell cancer Bi *et al.* study** [6]**, with a, percentage of mitochondrial counts among malignant and TME cells,**

**and b, dissociation stress among malignant and TME cells**. Dissociation stress is measured by the score of the meta-dissociation stress signature devised as the genes commonly found in all signatures of dissociation (Methods).

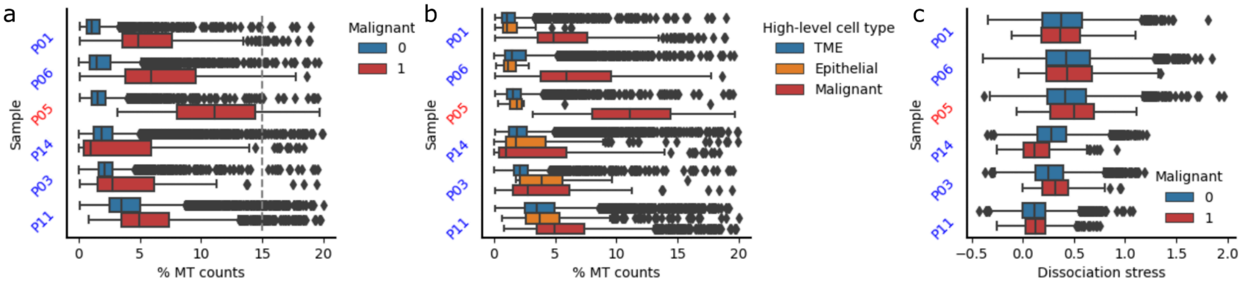


**Supplementary Figure S9: Distribution per patient of the nasopharyngeal carcinoma Chen *et al.* study** [3], **with a, percentage of mitochondrial counts among malignant and TME cells, b, percentage of mitochondrial counts with the separation of TME and normal epithelial cells, and c, dissociation stress among malignant and TME cells**. Dissociation stress is measured by the score of the meta-dissociation stress signature devised as the genes commonly found in all signatures of dissociation (Methods).

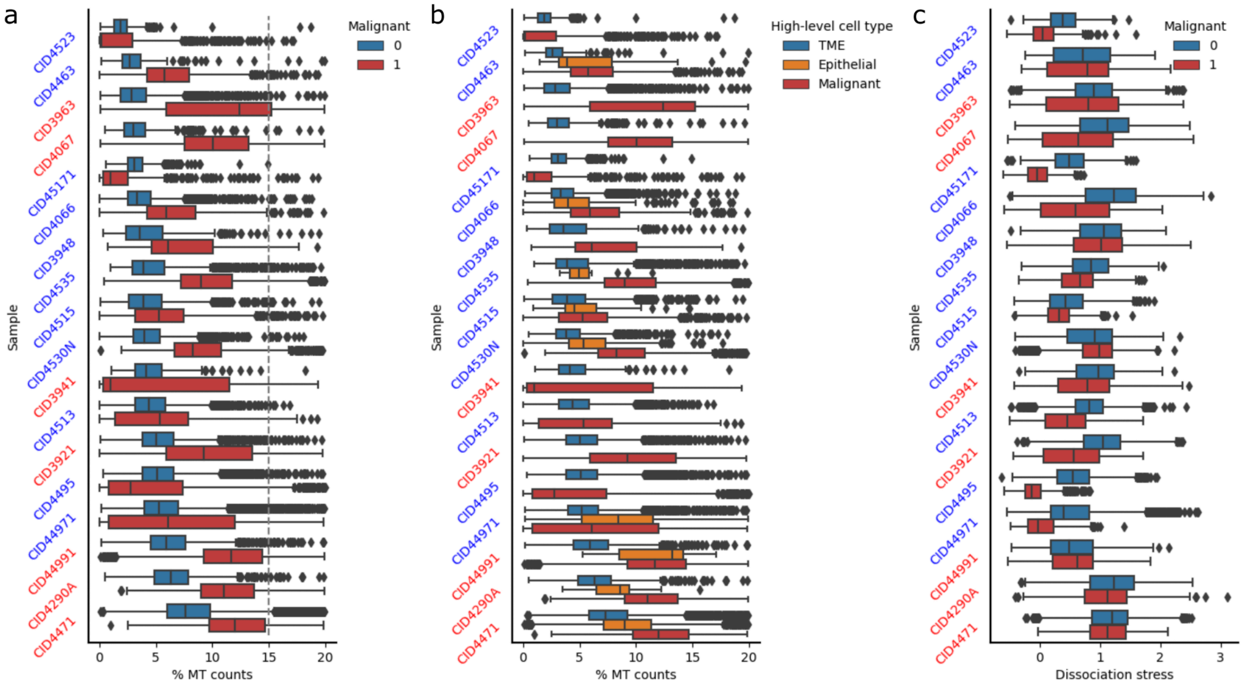


**Supplementary Figure S10: Distribution per patient of the breast cancer Wu *et al.* study** [7], **with a, percentage of mitochondrial counts among malignant and TME cells, b, percentage of mitochondrial counts with the separation of TME and normal epithelial cells, and c, dissociation stress among malignant and TME cells**. Dissociation stress is measured by the score of the meta-dissociation stress signature devised as the genes commonly found in all signatures of dissociation (Methods).

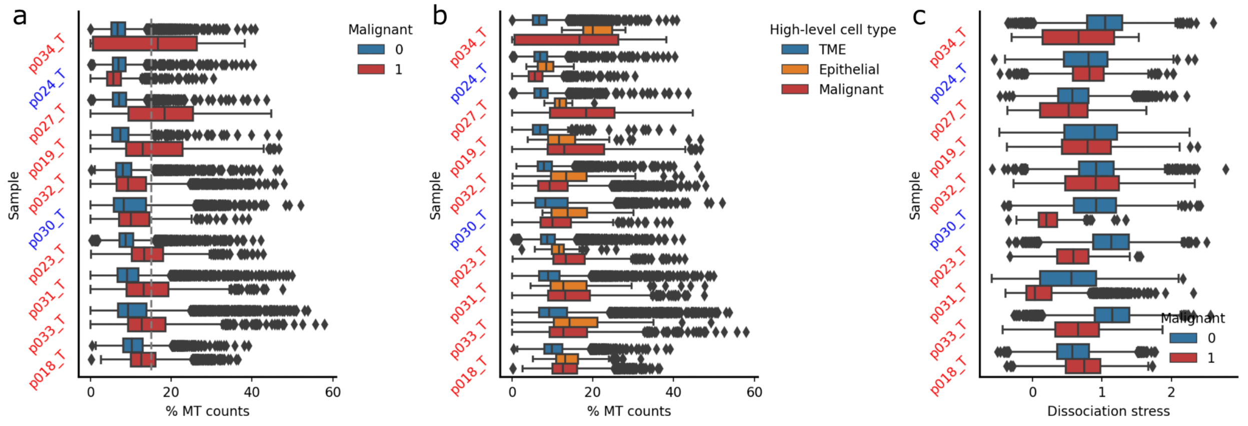


**Supplementary Figure S11: Distribution per patient of the lung adenocarcinoma Bischoff *et al.* study** [9]**, with a, percentage of mitochondrial counts among malignant and TME cells, b, percentage of mitochondrial counts with the separation of TME and normal epithelial cells, and c, dissociation stress among malignant and TME cells**. Dissociation stress is measured by the score of the meta-dissociation stress signature devised as the genes commonly found in all signatures of dissociation (Methods).

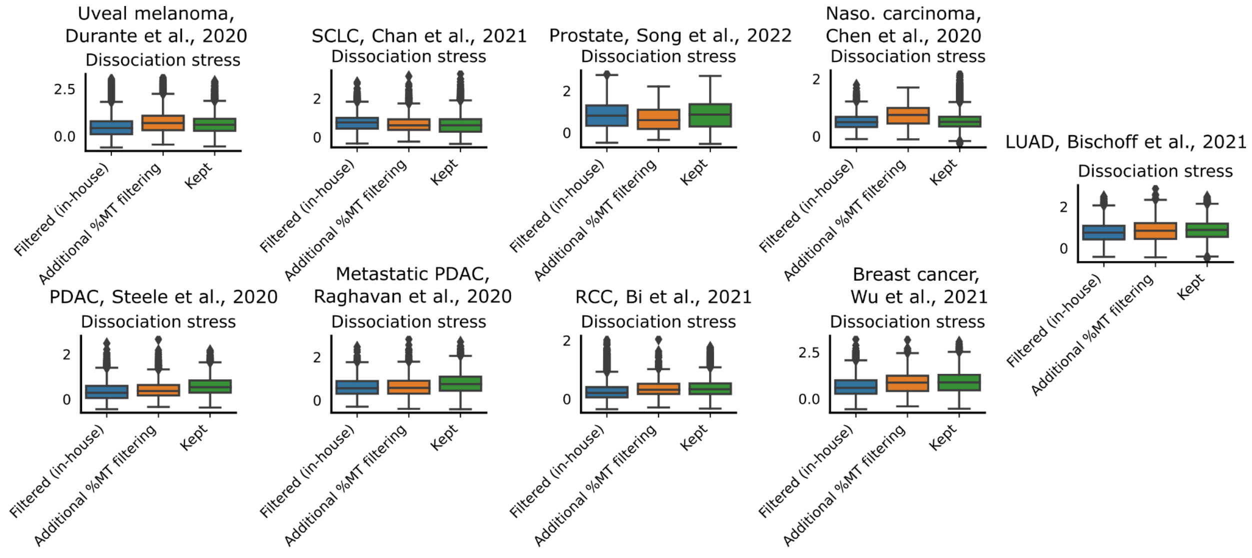


**Supplementary Figure S12: Dissociation-induced stress signature expression distribution in filtered and kept malignant cells.** For each study included in the analysis, we compute the dissociation stress for cells filtered using our QC procedure, cells kept using our procedure that present >15% MT counts, and all other remaining cells.

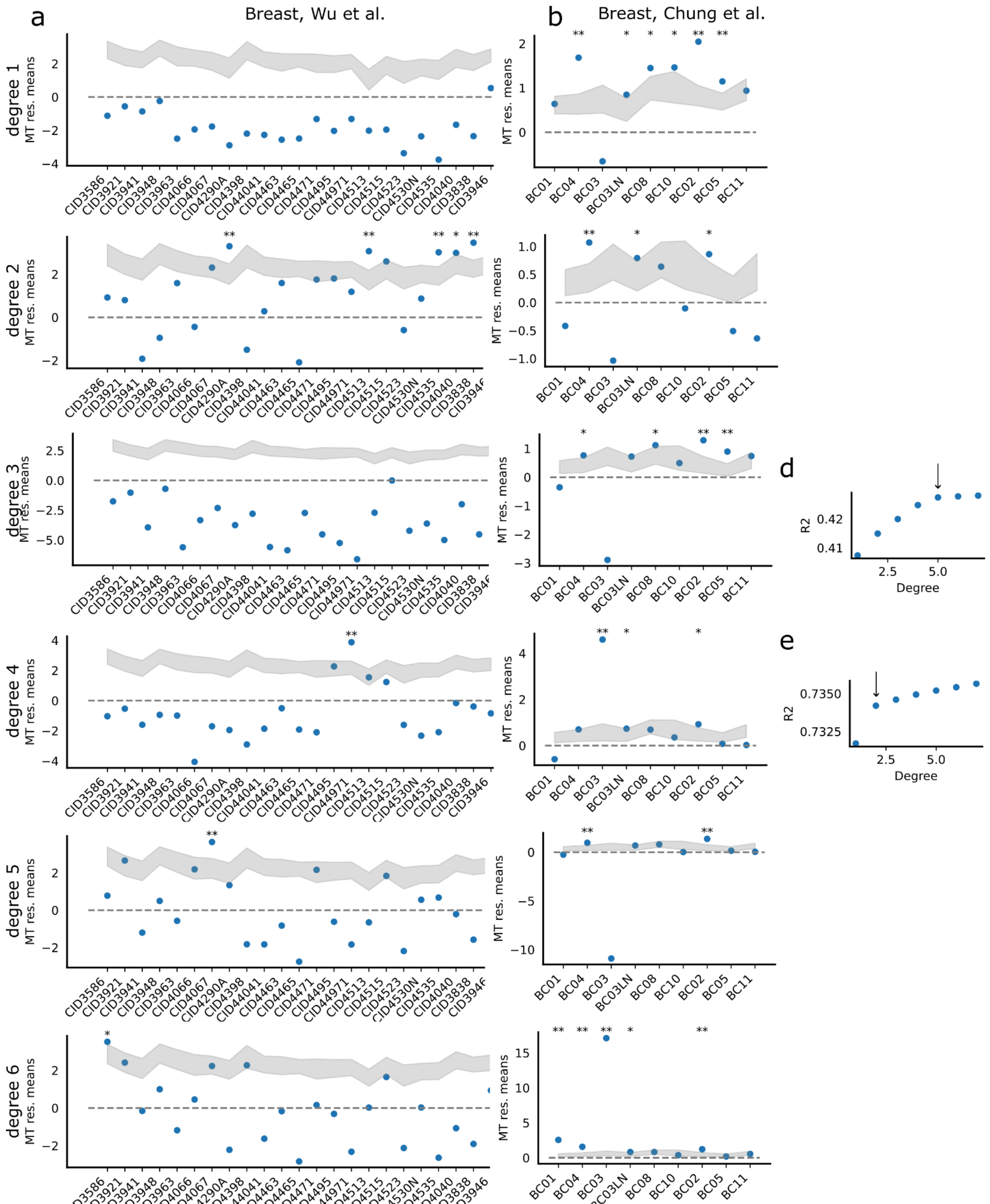


**Supplementary Figure S13: Residuals of MT-encoded genes for two breast cancer cohorts across different modelizations for a, the Wu et al.** [7] **and b, the Chung et al.** [13] **cohorts.** For each patient, paired bulk and single-cell measurements are available. We model the bulkified vs bulk relationship using a polynomial regression; we represent results for polynomial regression of degrees 1 through 6. The residuals represent the difference for each MT-encoded gene between the true and predicted bulkified values (Methods). To estimate whether a patient displays significantly differing residuals, we use an empirical one-sided test that tests the amount of times the mean residuals of the MT-encoded genes is higher than that of the randomly sampled genes with a similar level of expression. The 95% confidence interval of the mean residuals of randomly sampled genes is represented as a shaded gray area. **d-e,** Coefficient of determination (R2) for the different degrees for the **d,** Wu et *al.* and **e,** Chung et *al.* cohorts. The selected model is the one at the "elbow", i.e., when the increased complexity comes at the cost of minimal improvement of R2.

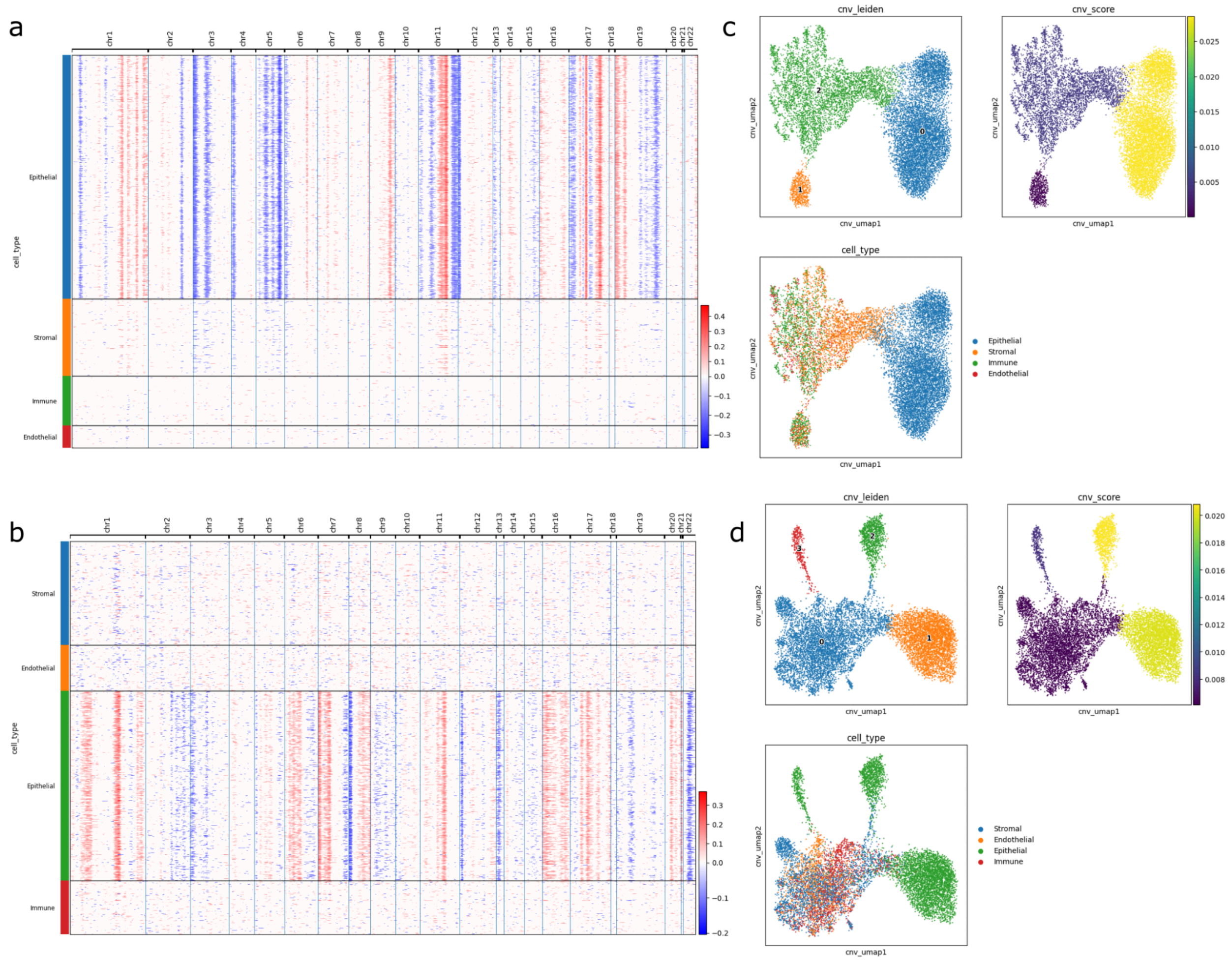


**Supplementary Figure S14**: **Copy number variation (CNV) analysis of metacells in breast ductal carcinoma in situ (DCIS) and lung adenocarcinoma (LUAD). a-b,** CNV profiles of metacells in **a,** DCIS and **b,** LUAD, with cell types assigned based on maximum canonical marker scores and inferCNV analysis using putative healthy cells as a reference. **c-d,** UMAP visualization of metacells with CNV data in **c,** DCIS and **d,** LUAD, colored by Leiden clustering, average CNV score, and putative cell type annotations. CNV clusters are categorized as malignant or healthy based on CNV scores, and cell type annotations are then adjusted accordingly.

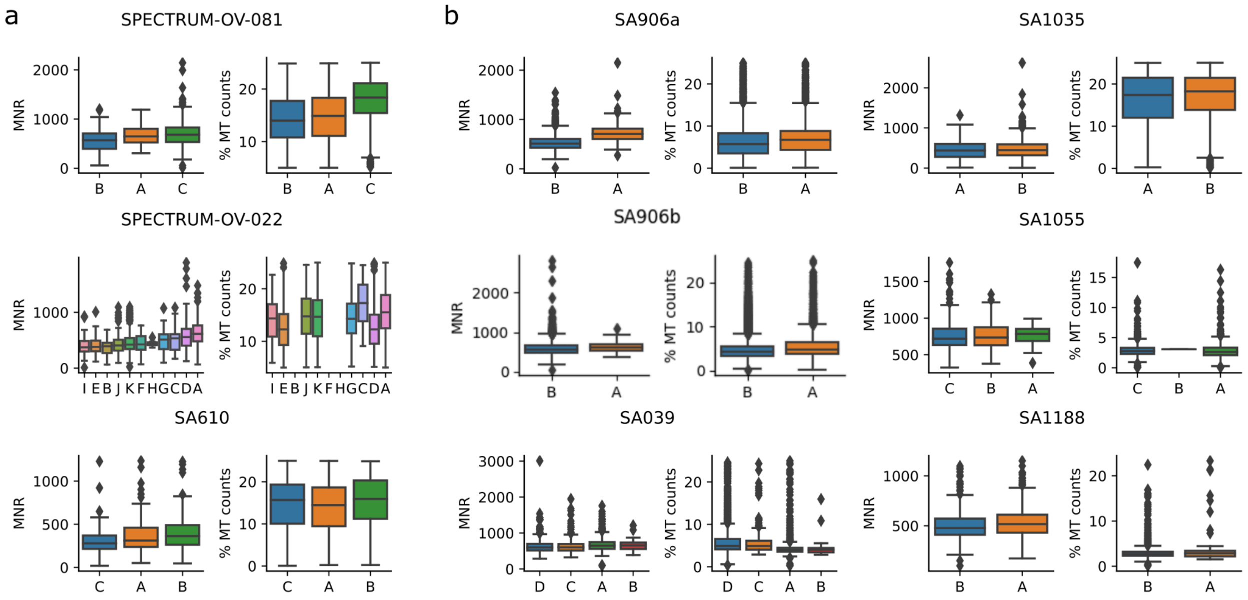


**Supplementary Figure S15:** Comparison of distribution between mitochondrial-to-nuclear gene ratio (MNR) and the percentage of mitochondrial counts (% MT counts) in a, ovarian cancer patient sample and b, engineered 184-hTERT cell lines from Kim et al. [14] Cells in scRNA-seq are computationally assigned to clones inferred in the DLP+-sequenced population using the TreeAlign algorithm [15]. The MNR is inferred using the genomic data while the % MT counts is computed using the scRNA-seq data. Clones with higher MNR often exhibit higher % MT counts.

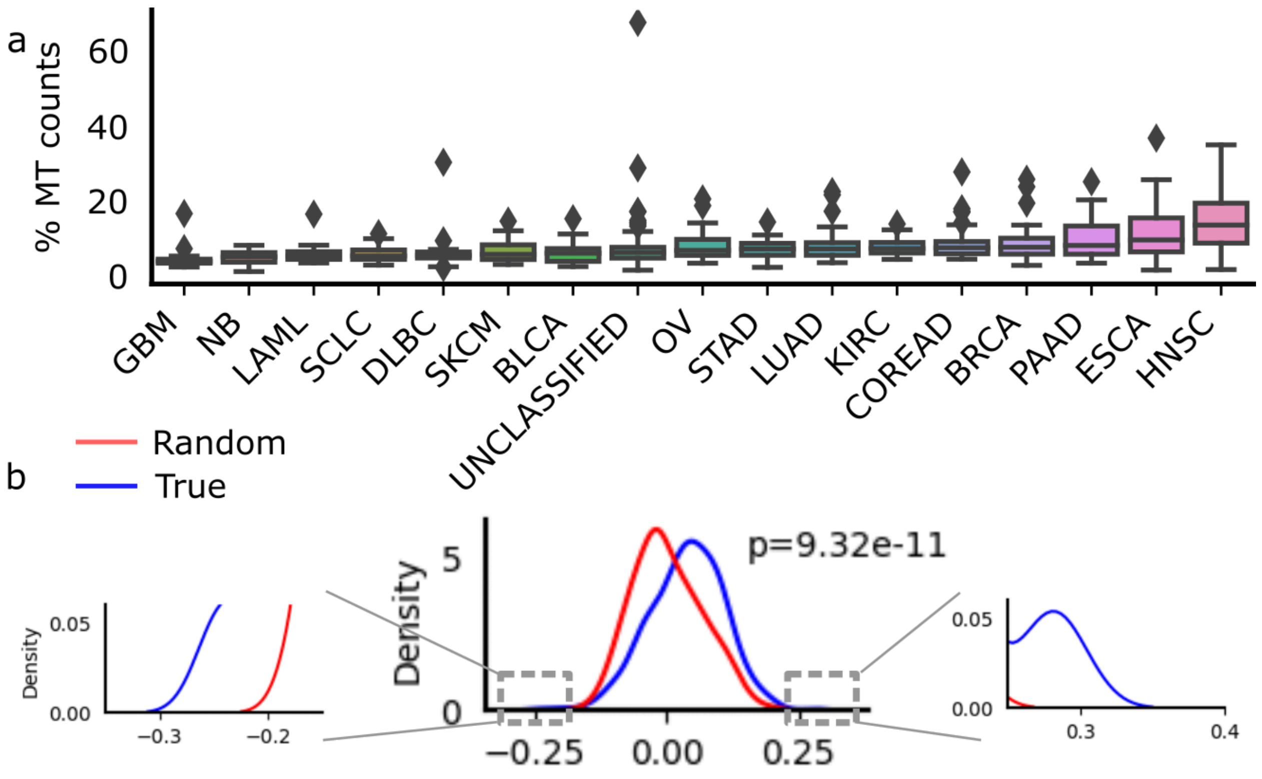


**Supplementary Figure S16: Drug resistance in cell lines. a,** Distribution of the % of mitochondrial counts in cell lines in the CCLE. **b,** Distribution of the median correlation across cell lines of specific cancer types between the pctMT and the IC50 of drugs. Significance of the difference between distributions is computed using a Kolgomorov-Smirnov test.

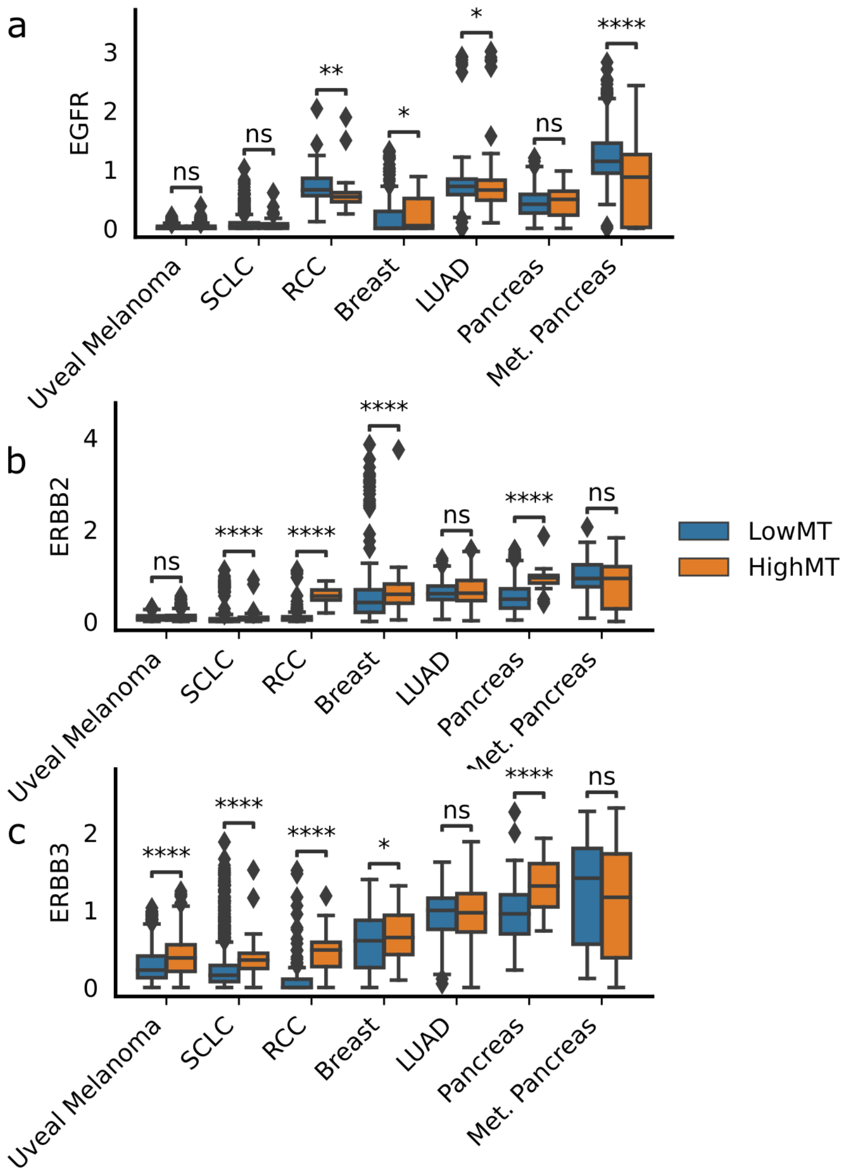


**Supplementary Figure S17: EGFR family gene expression across cancer types.** The log1p(CP10k) expression distribution is compared across HighMT and LowMT groups using a Mann-Whitney U test.

a

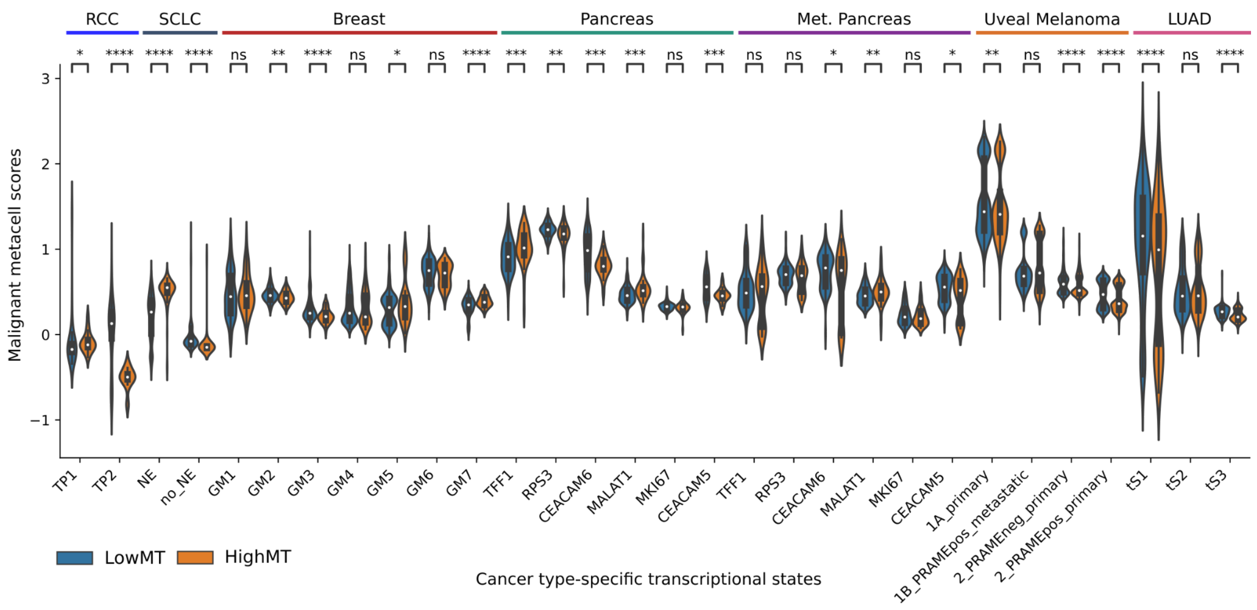


**Supplementary Figure S18: Association between pctMT and malignant cell states. a,** Distribution of scores of previously reported cancer-specific transcriptional states in LowMT and HighMT malignant cells. Significance is computed using Mann-Whitney U test on metacells. *: $0.01 \leq p < 0.05$; **: $0.001 \leq p < 0.01$; ***: $p < 0.001$.

| | Melanoma | SCLC | Pancreas | MetPancreas | RCC | Breast |
|---|---|---|---|---|---|---|
| **Median_High** | 0.554 | 0.713 | 0.584 | 0.313 | 0.774 | 0.666 |
| **Median_Low** | 0.585 | 0.714 | 0.593 | 0.249 | 0.574 | 0.68 |
| **Diff** | -0.031 | -0.001 | -0.009 | **0.064** | **0.2** | -0.015 |
| **p-value** | 0.0 | 0.357 | 0.13 | 0.001 | 0.0 | 0.163 |
| **q-value** | 0.0 | 0.391 | 0.177 | 0.004 | 0.0 | 0.255 |

**Supplementary Table 1: Expression of oxidative phosphorylation (OXPHOS) in the HighMT versus LowMT population.** OXPHOS is scored in metacells and the median for the HighMT and LowMT population, the difference in medians, and the p-value and FDR-corrected q-value for the difference computed with a Mann-Whitney U test are indicated for each cancer type.
